# Intracellular accumulation and secretion of hydrophobin-enriched vesicles aid the rapid sporulation of molds

**DOI:** 10.1101/2020.08.18.255406

**Authors:** Feng Cai, Zheng Zhao, Renwei Gao, Mingyue Ding, Siqi Jiang, Qi Gao, Komal Chenthamara, Marica Grujic, Zhifei Fu, Jian Zhang, Agnes Przylucka, Pingyong Xu, Günseli Bayram Akcapinar, Qirong Shen, Irina S. Druzhinina

**Affiliations:** The Key Laboratory of Plant Immunity, Jiangsu Provincial Key Lab of Solid Organic Waste Utilization, Nanjing Agricultural University, Nanjing, China; Fungal Genomics Laboratory (FungiG), Nanjing Agricultural University, Nanjing, China; Institute of Chemical, Environmental and Bioscience Engineering (ICEBE), TU Wien, Vienna, Austria; Key Laboratory of RNA Biology, Institute of Biophysics, Chinese Academy of Science, Beijing, China; Department of Medical Biotechnology, Institute of Health Sciences, Acibadem Mehmet Ali Aydinlar University, Istanbul, Turkey

**Keywords:** aerial hyphae, electron microscopy, extracellular vesicles, *in vivo* microscopy, intracellular vesicles, lipid bodies, *Pichia pastoris*, small secreted cysteine-rich proteins, *Trichoderma*, vacuoles, vacuolar multicisternal structures

## Abstract

Fungi can rapidly produce large amounts of spores suitable for aerial dispersal. The hydrophobicity of spores is provided by the unique amphiphilic and superior surface-active proteins – hydrophobins (HFBs) – that self-assemble at hydrophobic/hydrophilic interfaces and thus change surface properties. Using the HFB-enriched mold *Trichoderma* and the HFB-free yeast *Pichia pastoris*, we revealed a distinctive HFB secretory pathway that includes an intracellular accumulation of HFBs in lipid bodies (LBs) that can internalize in vacuoles. The resulting vacuolar multicisternal structures (VMS) are stabilized by HFB layers that line up on their surfaces. These HFB-enriched VMSs can move to the periplasm for secretion or become fused in large tonoplast-like organelles. The latter contributes to the maintenance of turgor pressure required for the erection of sporogenic structures and rapid HFB secretion by squeezing out periplasmic VMSs through the cell wall. Thus, HFBs are essential accessory proteins for the development of aerial hyphae and colony architecture.

## Introduction

The biology of filamentous fungi (mainly molds and mushrooms) is widely explained by the simple shape of their body, which is an apically growing branching tube or hypha. The physiochemical properties of the hyphal surface are important for fungal survival because fungi usually grow into their food or on the surface and feed by secreting digestive enzymes and subsequently absorbing dissolved small molecules. This nutritional strategy requires a large surface area/volume ratio (i.e., tubular shape) and a hydrophilic surface. However, fungi reproduce by spores that are not motile and therefore passively dispersed by air or water. Therefore, for reproduction and dispersal, fungi need to rapidly grow out of the substrate and form aerial and hydrophobic sporogenic structures (e.g., fruiting bodies, sporangia, and conidiophores) and spores^1, 2^ catching suitable dispersal conditions. The hydrophobicity of the spore or hyphal cell wall also influences their adhesion to the substrate and their interactions with symbiotic partners^3, 4^. Thus, the ability to modulate the hydrophobicity of the body surface and cross the hydrophilic/hydrophobic (i.e., water/air) interface is crucial for fungal lifestyle^5^.

One billion years of evolution of filamentous fungi^6^ has resulted in molecular adaptations to the physicochemical challenges associated with their lifestyle. For example, higher filamentous fungi commonly secrete hydrophobins (HFBs), which are small (usually < 20 kDa) amphiphilic and highly surface-active proteins^7, 8, 9^ that are characterized by the presence of eight cysteine (Cys) residues, and four of these residues form two Cys - Cys pairs. HFBs are initially secreted in a soluble form and then spontaneously localized at the hydrophilic/hydrophobic interface, where they assemble into amphipathic layers^10^ of varying solubility^7, 11^. These layers significantly decrease the interfacial tension, thus allowing the hyphae to breach the liquid surface and grow into the air. Fruiting bodies, aerial hyphae or spores are also largely coated by HFBs to reduce wetting, provide resilience to environmental stresses^2, 12^, promote the adhesion of spores and hyphae to hydrophobic surfaces or interactions with symbiotic partners^4^, and influence growth and development^13, 14, 15^.

Since fungi rapidly produce large amounts of spores^16^, they need to rapidly secrete large amounts of HFBs that need to be synthesized by sporulating aerial hyphae. In molds, aerial hyphae are ephemeral structures^5, 17^ that quickly become vulnerable and aged because they are deprived of nutrient and water absorption. In addition, aerial hyphae are frequently exposed to drought, UV radiation, and mechanical damage by rain, wind, or animals, and they may also need to grow against gravity. The massive formation of sporangia and spores is an energy-consuming task^16^ that leaves little resource for the secretion of HFBs. Under such circumstances, the mechanisms by which HFB can be massively secreted and how this process coordinates with the rapid sporulation of the fungus, allowing these proteins to play multiple roles in fungal reproduction, are not known.

Most fungi manage the above requirements with the aid of only a few HFB-encoding genes (JGI Mycocosm, June 2020), although some extremophilic species (*Wallemia ichthyophaga*^18^) or mycorrhizal mushrooms^19^ have a rich arsenal of HFBs. In Ascomycota molds, the genomes of fungi from the order Hypocreales have an extreme variety of HFB-encoding genes^20^. Among them, the genus *Trichoderma* exhibits the highest number and diversity of HFBs, which constitute the main genomic hallmarks of these fungi^20, 21^. The number of HFBs in individual species can range from seven in *T. reesei* to 16 in *T. atroviride*^20^. Compared with HFBs in mushrooms and some airborne Ascomycota, *Trichoderma* HFBs do not form rodlets or amyloids and are soluble in organic solvents and detergents^7, 11, 22^. An ecological genetic study of *T. guizhouense* and *T. harzianum* revealed that HFB4 on the spore surface controls the preferential dispersal mode and spore survival^2^. However, to date, the biological role of the enrichment of HFBs in the *Trichoderma* genomes has not been understood^20^. In the present study, we investigated the localization secretion mechanisms and function of HFBs during the development of the two commonly occurring cosmopolitan sibling *Trichoderma* species *T. harzianum* and *T. guizhouense*^23, 24^. The effect of HFB production on the cell ultrastructure was also investigated in the methylotrophic yeast *Pichia pastoris*, which has no HFB-encoding genes^25^.

## Results

### At least four HFBs are involved in the development of *T. harzianum* and *T. guizhouense*

The genomes of *T. harzianum* CBS 226.95 (Th) and *T. guizhouense* NJAU 4742 (Tg) contain eleven and nine HFB-encoding genes, respectively^2^. An expression analysis performed at the four stages of the life cycle (Supplementary material 1 Table S1.1) revealed that in both species, the highest transcription level of *hfb* genes was recorded during the formation of aerial hyphae. A principal component analysis (PCA) of the expression profile of *hfb* genes confirmed the strong involvement of *hfb4* and *hfb10* in the development of Tg and Th and a minor role of *hfb2* (Supplementary material 1 Figure S1.1). The remaining genes were hardly expressed at all stages except *hfb3*, which was only noticeable at the aerial hyphal formation stage. Thus, we constructed a library of *hfb*-deficient, hfb-overexpressing and fluorescently labeled mutants for the above four genes (Supplementary material 2 Table S2.1). As fluorescent protein tags may potentially influence the properties of the fused proteins with HFBs^26^, we first predicted properties of the fusion proteins using *in silico* 3D modeling and molecular dynamic (MD) simulation analysis for their behavior in water and then designed such fusion constructs that correspond to the highest probability of the exposed position of the HFB hydrophobic patch (Supplementary material 3). To verify that none of the observed differences were caused by genetic transformation, each genotype was represented by multiple mutants. We also produced reverse complemented mutants in which fluorescently tagged and untagged HFB-encoding genes were reintroduced to the corresponding deletion mutant and compared their properties (Supplementary material 4).

The deletion of either *hfb4* or *hfb10* or both genes resulted in reduced conidiation and a “wetted hyphae” phenotype (reduced surface hydrophobicity) in both species^2^. The deletion of *hfb2* did not result in any phenotypic alterations (see detailed phenotypic characterization in Supplementary material 5).

### HFBs massively accumulate in aerial hyphae of filamentous fungi and yeasts

*In vivo* epifluorescence microscopy of mutant strains expressing fluorescently labeled HFBs revealed that the formation of aerial hyphae was accompanied by massive intracellular accumulations of HFB4 and HFB10, which were stored in different types of membrane-bound vesicle-like organelles and in the periplasm (Fig. 1). HFB3 also showed intracellular accumulation. However, as this protein was more visible on the surface and at collarettes of sporogenic cells on conidiophores (phialides) (Fig. 1) and had very low expression levels, it was not included in the subsequent study. HFB4 and HFB10 were not associated with the cell wall or present in the cytoplasm. Even though both proteins were detected intra- and extracellular, HFB4 has affinity for stalk cells and phialides (Fig. 1, Supplementary materials 6 Figure S6.1). In contrast, HFB10 had a higher affinity than HFB4 to solid/liquid interfaces on the microscopy glass slide outside the cells.

**Figure 1.**
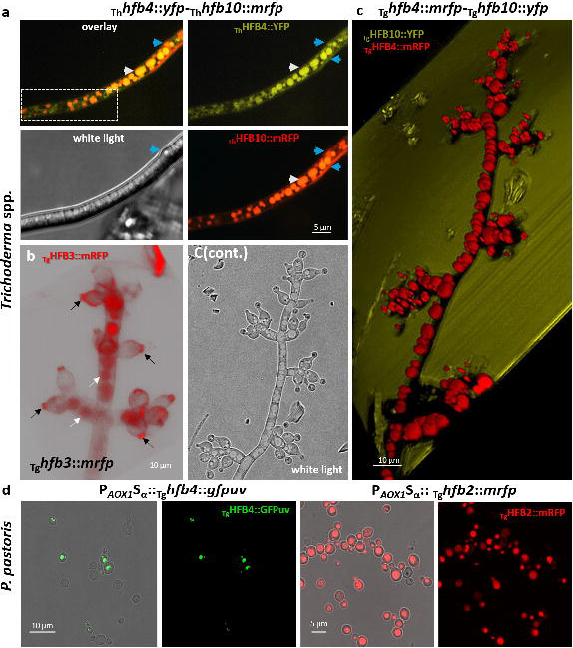
*In vivo* visualization of intracellular accumulation of fluorescently labeled hydrophobins in *Trichoderma* spp. and *Pichia pastoris* yeast. **a,** Aerial hyphae of *T. harzianum* _Th_*hfb4::yfp-hfb10-.:mrfp* accumulating HFBs in organelles, including putative conventional and tubular vacuoles (dashed area), putative vacuoles (white arrows) and the periplasm (blue arrow). **b,** Localization of _Tg_HFB3::mRFP in vacuole-resembling organelles (white arrows) and on the surface of phialides near the collarette (black arrows). **c,** 3D reconstructions of fluorescent overlay images of the mature conidiophore of *T. guizhouense* producing HFB4::mRFP and HFB10::YFP. The white light image of the same conidiophore is shown on the separate panel (**c**_continued_). **d**, *In vivo* confocal microscopy *of P. pastoris* producing fluorescently labeled HFB4::GFPuv (left) and HFB2::mRFP (right) from *Trichoderma* using the signal peptide of *S. cerevisiae* α-mating factor (Sα) under the control of the AOX1 promoter (P_*AOX1*_). HFB production was induced by methanol in *P. pastoris.*

The enrichment of intracellular HFB4 and HFB10 in conidiophores indicated that their intracellular localization was likely not linked to the potential recognition of a fusion construct as an alien protein. Nevertheless, to test whether the accumulation of HFBs in vesicle-like organelles were linked to the presence of the fluorescent tag, we expressed mRFP (without HFBs) using the promoter and signal peptide of _Tg_HFB4 (_Tg_P_*hfb*4_::mrfp) (Supplementary materials 7). The resulting protein was secreted into the medium without any sign of intracellular localization (Supplementary material 7 Figure S7.1). To test whether the intracellular accumulation of HFBs would occur in other fungi, we overexpressed *Trichoderma* HFB-encoding genes in *P. pastoris* (which has no *hfb* genes in its genome^25^) (Supplementary material 2 Table S2.1). Despite the presence of the signal prepropeptide of the α-mating factor from *Saccharomyces cerevisiae, P. pastoris* cells also massively accumulated HFBs intracellularly (Fig. 1 d). Together, these results indicate that the intracellular localization of fluorescently tagged HFBs in vesicle-like organelles and in the periplasm is not led by the signal peptide or due to the tag but linked to the properties of HFBs.

The morphology of HFB-containing organelles closely resembled vacuoles and lipid bodies. To test for the vacuolar location of HFBs, we deleted the gene encoding endosomal RAB7 (OPB37336), which is a small GTPase required for the late steps of multiple vacuole delivery pathways^27^ (homologs include *ypt7*^28^ in *Saccharomyces cerevisiae* and *avaA* in *Aspergillus nidulans*^27^) in Tg and Tg mutants expressing HFB4::mRFP *(_Tg_hfb4::mrfp* strain). The mutants *Δrab7* and _Tg_*hfb4*::*mrfp*-_Tg_Δ*rab7* showed strongly reduced growth with abolished ability to form aerial hyphae, were highly hydrophilic, did not produce conidiophores or conidia and had no vacuoles (Fig. 2, Supplementary material 8 Figures S8.1 – S8.2). The cytoplasm of the _Tg_*hfb4*::*mrfp*-_Tg_Δ*rab7* strain was enriched in small HFB4::mRFP-containing vesicles (heterogeneous appearance of cytoplasm) but lacked large HFB-containing organelles. Thus, these data confirmed the vacuolar localization of HFBs.

**Figure 2.**
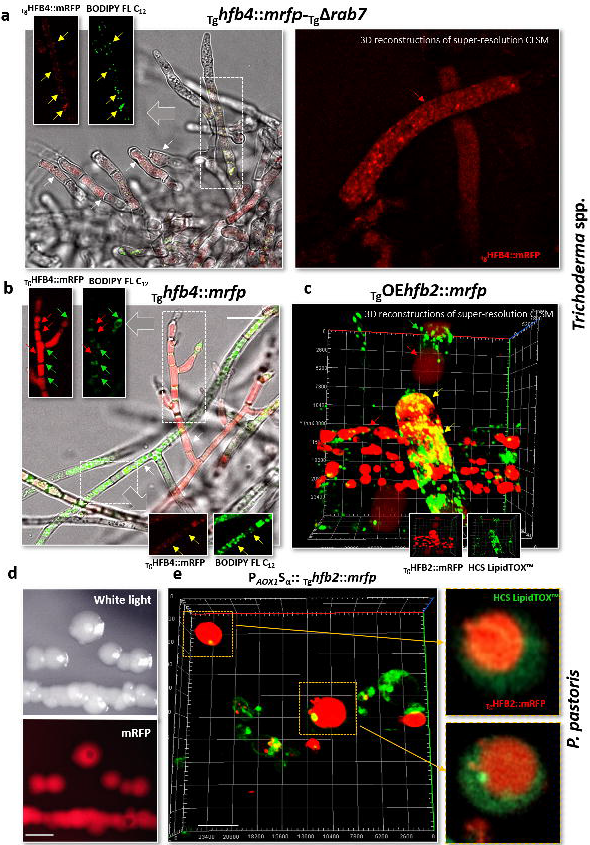
Accumulation of HFBs in vacuoles and lipid bodies. **a,** Fluorescent overlay images of the phospholipid-specific stained (using a green fluorescent fatty acid (BODIPY™ FL C_12_)) aerial hyphae of the *Trichoderma* mutant producing HFB4::mRFP but lacking the key protein for the cytoplasm-to-vacuole sorting pathway, RAB7 (homologous to YPT7 in *S. cerevisiae)*, and 3D reconstructions of the corresponding hyphae. **b,** Fluorescent overlay images of the phospholipid-specific stained (using BODIPY™ FL C_12_) aerial hyphae of the Tg mutant producing _Tg_HFB4::mRFP. **c,** 3D reconstructions of phospholipid-specific stained (using HCS LipidTOX™ Green) aerial hyphae of the mutant overproducing HFB2::mRFP. Scale bar= 10 μm**. d,** Stereo confocal microscopy *of P. pastoris* producing fluorescently labeled HFB2::mRFP from *Trichoderma* using the signal peptide of *S. cerevisiae* α-mating factor (Sα) under the control of the AOX1 promoter (P_*AOX1*_). HFB production in *P. pastoris* was induced by methanol. Scale bar = 5 mm. **e,** Intracellular accumulation of HFB2::mRFP in the same strains as in **d**. Scale bar = 10 μm. Lipids were stained using HCS LipidTOX™ Green. The inserts in **a** – **d** correspond to framed areas showing the localization of the fluorescently labeled HFBs (red arrows) and phospholipids (green arrows) or both (yellow arrows).

To test whether the other type of HFB-containing organelles observed in Fig. 1 corresponded to lipid bodies, we applied differential phospholipid stains including BODIPY™ FL C_12_ and HCS LipidTOX™ Green reagent (Thermo Fischer Scientific, USA) to the _Tg_*hfb4::mrfp* and _Tg_*hfb4::mrfp*-_Tg_Δrab7 strains (Fig. 2). These results indicated that in lipid-rich hyphae lacking large vacuoles, HFB4 was localized in lipid bodies, while in vacuole-rich cells, such as aerial hyphae and conidiophores, HFB4 preferentially accumulated in vacuoles (Fig. 2). In the absence of vacuoles in the *Δrab7* strain, HFB4 could be detected in small vesicles and lipid bodies (Fig. 2 b). Similar results were obtained for HFB2. In agreement with the results on *Trichoderma*, in *P. pastoris*, intracellular fluorescently tagged HFBs were localized in lipid bodies and vacuole-like organelles (Fig. 2 c).

For transmission electron microscopy (TEM) analysis of cell ultrastructure, we constructed strains constitutively producing a fluorescently tagged HFB to secure a homogenous high expression level of HFB in the sample. For this purpose, we selected a phenotypically neutral gene *hfb2.* The resulting mutants (_Tg_OE*hfb2*::*mrfp*) appeared visibly pink (Supplementary material 9) accumulating large amounts of the fusion proteins intracellularly. It also revealed that aerial hyphae can massively accumulate and tolerate a large quantity of HFBs (Fig. 3 a; see the corresponding video in Supplementary material 10). TEM analysis of the aerial hyphae of the parental and overexpressing strains compared with those of the deletion strains revealed numerous vacuolar multicisternal structures (VMSs) associated with the high production of HFBs (Fig. 3 b). VMSs are characterized by a “matryoshka” of electron-dense membranes or films, and this result was found with the three HFBs studied in both species (Fig. 3). The matrix of the VMSs was almost electron-transparent, with frequent inclusions of various sizes and electron densities (Fig. 3). The lipid bodies in HFB-overproducing cells were relatively more electron dense than those in the cells of the wild type and strains lacking HFBs, suggesting differences in their chemical composition such as putative protein enrichment (*see above).* Multiple examples of lipid bodies internalizing in vacuoles (Fig. 3) suggested that this process could be important for VMS and “matryoshka” formation. The hyphae of mutants lacking HFBs (in particular, _Tg_Δ*hfb4*-_Tg_*Δhfb10* or _Th_*Δhfb4-_Th_Δhfb10*, Fig. 3 b) contained lipid bodies and vacuoles but were deprived of VMS. Interestingly, the periplasm volume appeared to be essentially smaller in mutants lacking HFB4 and HFB10 than in the parental and HFB-overexpressing strains (Fig. 3 b). At several sites, the release of VMS content into the periplasmic space was imaged (Fig. 3, also see below), indicating that HFB secretion may go through the initial localization of HFB-enriched vesicles such as VMS in periplasm.

**Figure 3.**
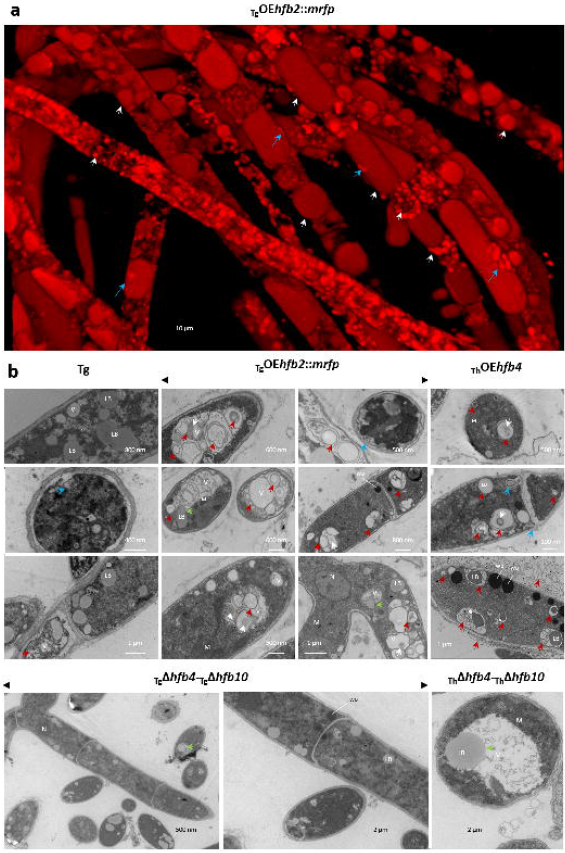
Vacuolar multicisternal structures (VMSs) in *Trichoderma* hyphae producing HFBs. **a**, Young aerial hyphae of *T. guizhouense* producing HFB2::mRFP under the control of the strong constitutive promoter of the *cdna1* gene from *T. reesei* QM6a. White and gray arrows point to the different types of organelles or vesicles accumulating HFB2::mRFP. The blue arrow indicates a putative site of organelle rupture or periplasmic accumulation. Supplementary material 10 contains a video corresponding to this image. **b,** Representative TEM micrographs showing the hyphal ultrastructure of strains including the wild-type strain of *T. guizhouense* (Tg), HFB2::mRFP-overexpressing strains of Tg (as on **a**), HFB4-overexpressing *T. harzianum* (Th), or hyphae of Tg and Th lacking HFB4 and HFB10. All images of the aerial hyphae were obtained from a two-day-old colony. V – vacuole, LB – lipid body, N – nucleus, M – mitochondrion, WB – Woronin body, and pP – putative protein inclusion. Red arrows point to putative HFB vesicles in vacuolar multicisternal structures (VMSs) and “matryoshka” structures. Blue arrows indicate the release of VMS content to the periplasm (see **a**). Green arrows highlight the internalization events of LBs to Vs. Representative images were selected from a total of 695 images obtained for Tg (361) and Th (334). Samples for TEM were prepared with at least two mutants and 30 images studied per genotype. Mutants are listed in Supplementary material Table S2.1.

The ultrastructure of HFB-overproducing *P. pastoris* cells resembled that of *Trichoderma* with abundant VMS having the characteristic “matryoshka” appearances except that VMSs were not linked to the periplasm (Supplementary material 11 Figure S11.1). Similar to *Trichoderma*, multiple events of lipid bodies internalized in vacuoles were also observed in *P. pastoris* (Supplementary material 10 Figure S11. 2). In this yeast, the release of HFBs occurred by the bursting of cells overloaded with intracellular HFBs.

Because VMS membranes can originate from the endoplasmic reticulum (ER) as ER whorls^29^ and the massive accumulation of membranous structures in vacuoles resembles autophagy morphology^30, 31^, we tested the expression of autophagy-related genes. The results showed no significant upregulation of autophagy marker genes associated with high *hfb* gene activity (aerial hyphae) (Supplementary material 12 Figure S12.1). Compared to *Trichoderma*, the accumulation of HFB-enriched VMSs did not cause any considerable upregulation of autophagy marker genes in *P. pastoris* (Supplementary material 12 Figure S12.1).

Taken together, the results of the above analyses indicated that HFBs first accumulated in lipid bodies that were internalized in vacuoles and likely gave rise to VMSs. In VMSs, HFBs self-assembled in the hydrophobic/hydrophilic interface led to the lining up of VMS membranes and/or formation of HFB vesicles similar to the bilayer HFB1 vesicles reported by Hähl et al. in a cell-free system^32^. In some cells, several VMSs containing HFB vesicles formed gigantic central tonoplasts, as observed in Fig. 3 a. In aerial hyphae, the secretion of HFBs occurred through the release of VMSs with visually stable HFB microvesicles to the periplasm (Fig. 3; *see below*). The observed affinity of HFBs for lipid bodies and VMSs was caused by the core HFB sequences and was not influenced by the signal peptide or fluorescent tags, as indicated by the expression patterns in the respective *T. guizhouense* and *P. pastoris* mutants.

### HFBs aid conidiophore erection by contributing to turgor pressure in aerial hyphae

The deletion and overexpression of HFBs influenced the structure of mature aerial hyphae (Fig. 4). A cryo-scanning electron microscopy (cryo-SEM) analysis of strains lacking either HFB4 or HFB10 or both revealed multiple large indentations (0.3 – 1 μm in diameter) on the surface of aerial hyphae, including on the hyphal tips (Fig. 4 a), where the turgor pressure is the highest^33^. In contrast, HFB-overexpressing strains had a visually “swollen” appearance (Fig. 4 b and c). Moreover, conidiophores and aerial hyphae of the HFB-overexpressing and wild-type strains of both species had abundant herniations (protrusions) of 0.3 to 1.3 μm in diameter (Fig. 4) filled with HFB vesicles.

**Figure 4.**
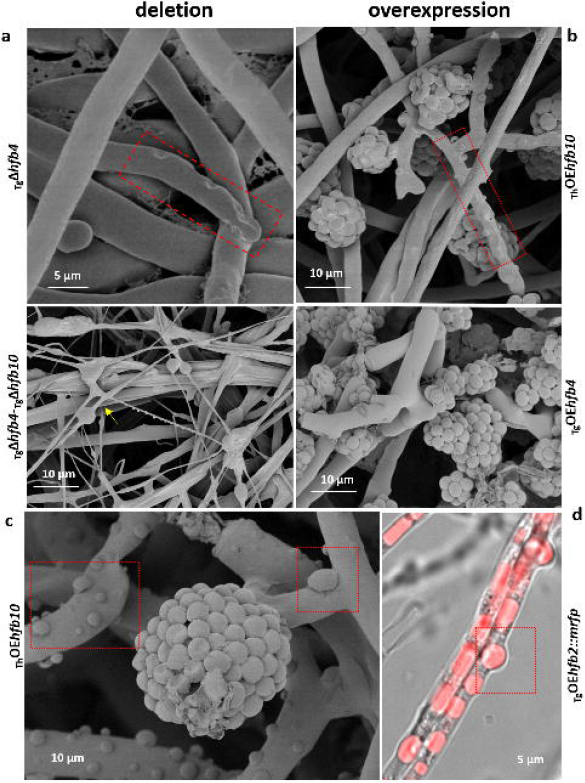
Turgor pressure in aerial hyphae of HFB-deficient and HFB-overexpressing mutants of *Trichoderma* spp. **a,** Cryo-SEM micrographs of aerial mycelium of HFB-deficient *Trichoderma* cultures grown on PDA for 11 d. Yellow arrows point to remains of burst hyphae. **b-c,** Cryo-SEM micrographs of aerial mycelium of HFB-overexpressing *Trichoderma* cultures grown on PDA for 28 d. **d,** An image from epifluorescence microscopy showing the HFB-rich content of protrusions corresponding to those shown in **b** and **c**. Red boxes in **a** – **d** highlight indentations or herniations corresponding to putatively reduced or increased intracellular turgor pressure, respectively. Representative images were selected from a total of 523 images obtained for Tg and Th. Samples for SEM were prepared with at least two mutants and 30 images studied per genotype. Mutants are listed in Supplementary material Table S2.1.

These phenotypes and the multiple observed cases of nonapical cell wall rupture, hyphal herniation and bursting (Fig. 4 and Fig. 3) suggest that HFBs are linked to the regulation of turgor pressure in aerial hyphae. In such hyphae, which are hydrophobic and therefore deprived of water influx and less influenced by osmosis, turgor pressure can be maintained by the formation of a central tonoplast^34^. HFB vesicles can contribute to the formation and mechanical stability of such organelles.

Turgor pressure is the internal force that pushes the cytoplasm against the cell wall^33^. Thus, the indentations and herniations observed in HFB mutants could develop due to changes in the cell wall structure, as HFBs were reported to affect this organelle^35^. However, analysis of the cell wall ultrastructure did not reveal any visible effect of HFBs on the hyphal cell wall (Supplementary material 13 Figure S13. 1) except that older hyphae of parental or HFB-overexpressing strains had a thick extracellular matrix that does not explain the observed herniations. The reduced abundance of conidia observed in both *hfb*-lacking and *hfb*-overexpressing strains (Supplementary material 5 Figure S5.2-5.4) also supported the hypothesis that a disorder in HFB content altered conidiophore formation because of insufficient turgor pressure for conidiophore uprising and the requirement of this step for subsequent conidiation. Similarly, the reduced rate of surface growth observed for Tg strains lacking the *hfb4* gene (Supplementary material 5 Figures S5.2) could also be linked to impaired turgor pressure, which is required for apical hyphal growth^33^. Considering the proposed mechanism for HFB secretion by aerial hyphae through the initial accumulation of HFB-enriched vesicles in the periplasm, elevated turgor pressure was probably also required for the facilitated secretion of HFBs by squeezing the periplasm space due to the intracellular force on the plasma membrane against the cell wall.

Aerial hyphae are ephemeral. Therefore, on the eleventh day of cultivation, the aerial parts of the colonies of both species were partially degraded, and vital hyphae were maintained only around conidiophores (Fig. 4). At this stage, the strains lacking the major HFBs had almost no intact aerial hyphae because most of them had already burst, resulting in a “spaghetti with mozzarella” SEM phenotype (Fig. 4 a), where regular hyphae were intermixed with thin hyphal debris (putatively empty cell walls).

We thus speculate that HFB-lined VMSs, which may be either single HFB vesicles or large aggregates of such vesicles, contribute to the development of turgor pressure in aerial hyphae, including conidiophores, in a manner similar to that of tonoplasts in plant cells^36^. This hypothesis is also supported by the observations of enlarged bursting cells of *P. pastoris* overexpressing HFBs. We propose that this fungus has no mechanisms for targeting HFB vesicles to the periplasm; therefore, excess HFBs result in cell rupture due to increased intracellular pressure. Furthermore, the lack of conidiophores and generally scarce aerial mycelium of *Δrab7* mutants that are unable to form vacuoles, VMS and tonoplasts can also be explained by insufficient turgor for conidiophore formation.

### Aerial hyphae secrete HFBs for the formation of a massive extracellular matrix

The massive accumulation of HFBs in aerial hyphae is visible without magnification if fluorescently labeled mutants are placed under a fluorescent imaging system, such as ChemiDoc MP (Bio-Rad, USA) (Fig. 5, Supplementary material 14). The time course observation showed the coordinated appearance of HFB4 and HFB10 prior to conidiation along with the formation of conidiophores (Supplementary material 15). Monitoring of *hfb* gene expression during conidiogenesis demonstrated that at least four *hfb* genes *(hfb2, hfb3, hfb4*, and *hfb10)* were significantly (up to several thousand-fold) upregulated precisely at the beginning of conidiophore formation compared to hyphal growth. In both species, the expression of *hfb3* was clearly apparent at this stage, while it was not considered significant when the entire aerial mycelium was tested (Supplementary material 1). Observation of the colony surface using confocal microscopy revealed an HFB-enriched protein matrix surrounding the spores and conidiophores (Fig. 6 a). A 3D reconstruction of the colony surface showed that the matrix has an uneven distribution of HFB4 and HFB10 (see the video of 3D reconstruction in Supplementary material 15), with HFB4 being more associated with the spores and HFB10 with the hyphae.

**Figure 5.**
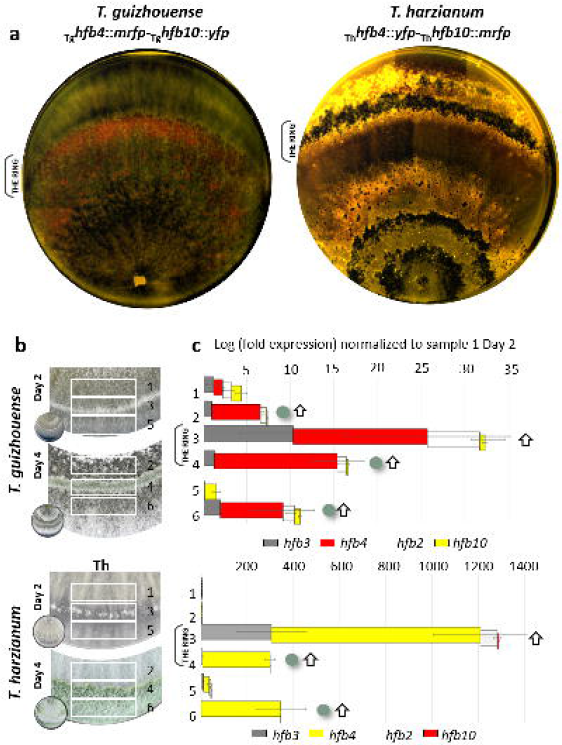
Colony architecture and dynamic HFB accumulation in aerial hyphae during conidiogenesis. **a,** Fluorescent imaging of conidiating *Trichoderma* colonies with labeled HFB4 and HFB10. The time course images based on a similar principle are shown in Supplementary material 14. Note that HFB4 is labeled with red and yellow fluorescent proteins in Tg and Th, respectively, while HFB10 is labeled with yellow and red fluorescent proteins in Tg and Th, respectively. **b,** Formation of the conidiating ring during the fine time course of conidiogenesis of *Trichoderma* spp. Photos show sampled areas of *Trichoderma* cultures for gene expression analysis (boxed). **c,** Relative expression of *hfbs* during the fine time course of conidiogenesis of *Trichoderma* spp., quantified by qPCR. The values are normalized to those of the housekeeping gene *tef1* and expressed in relation to aerial hyphae prior to conidiogenesis (position 1 in **b**). Horizontal bars indicate standard deviations. Green dots and white arrows indicate spores and conidiophores appearing in stages, respectively.

**Figure 6.**
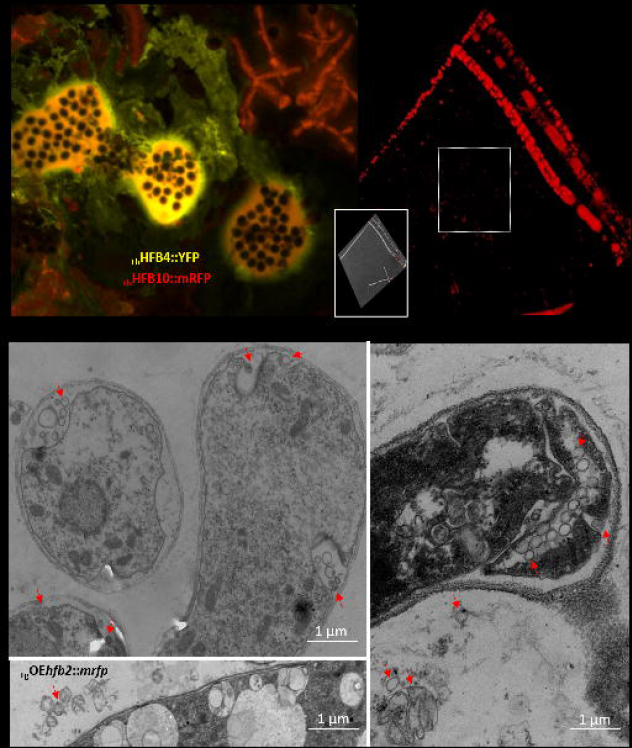
Extracellular HFB matrix, extracellular HFB-enriched vesicles and periplasmic vesicles. **a,** A confocal microscopy image of the print of a conidiating colony of *T. harzianum _Th_hfb4::yfp-_Th_hfb10::mrfp* on a cover glass without the addition of extra water. The images of the corresponding plates are shown in Supplementary material Figure S5.1. Supplementary material 15 contains 3D reconstructions of such matrices for both species. **b,** Intra- and extracellular HFB-enriched vesicles washed out from the *T. guizhouense* culture overexpressing HFB2::mRFP (an overlay image is shown in the inset). Supplementary material 16 contains videos recording these vesicles using an epifluorescence microscope. Proteomic characterization of these vesicles is given in Supplementary material 9. **c**, TEM micrographs of 48-h-old *Trichoderma* spp. hyphae overexpressing HFBs without and with fluorescent tags. Red arrows indicate vesicles that may correspond to those seen in **b**. Representative images were selected from 523 images obtained for Tg. Samples for TEM were prepared with at least two mutants and 30 images studied per genotype. Mutants are listed in Supplementary material Table S2.1.

These results are concordant with the hypothesis that HFBs facilitate the formation of aerial hyphae and conidiophores (through increased intracellular turgor) and thus modulate the colony architecture. The analysis of mycelial wash of _Tg_*OEhfb2::mrfp* revealed extracellular vesicles (Fig. 6 b; Supplementary material 16) that were enriched in HFBs but also contained other proteins (Supplementary material 9 Figure S9.1 d). The TEM micrographs of these hyphae also showed extracellular vesicles. We propose that the massive secretion of HFBs in aerial hyphae follows their accumulation in the interface of intracellular vesicles such as VMS. HFB-enriched vesicles are then released through the plasma membrane to the periplasm, where they are squeezed out through the cell wall and partially remain intact. The release of large VMS may cause hyphal bursts (Fig. 4). The process is further aided by the increased intracellular turgor pressure. Thus, we propose that the massive secretion of HFBs by aerial hyphae is a prerequisite for conidiation not only because HFBs are essential spore protectants^2, 10^ but also because they aid the formation of all hydrophobic structures of fungal colonies, such as aerial hyphae, conidiophores, and spores.

## Discussion

Since their discovery, HFBs have been known to play multiple extracellular roles in filamentous fungi^15, 37^. Located outside of the cell, HFBs assemble at hydrophobic/hydrophilic interfaces, modulate surface properties, and mediate a broad spectrum of fungal interactions with their environment^8^. In molds (Ascomycota), HFBs coat spores, thereby protecting against abiotic stresses^2, 38, 39^ and supporting attachment to hosts or substrates^40^. Because molds can rapidly generate large amounts of spores^16^, HFBs must also be abundantly secreted. However, attempts to produce these proteins in filamentous hosts using standard fermentation protocols have proven to be inefficient^41, 42^ likely because the submerged hyphae need their surface to be hydrophilic for nutrient acquisition, which can be compromised by the access of HFBs.

Here, we uncovered a distinctive secretory pathway of HFBs in aerial (nonfeeding and hydrophobic) hyphae that includes the formation of putative HFB-enriched vesicles^32^ for an intracellular accumulation step that is followed by transport to the periplasm and release to the outer space by squeezing through the cell wall. Our results show that this mechanism is tightly linked to HFB properties and is strictly functionally determined. Consistent with previous studies, we also detected that the high expression of HFB-encoding genes is characteristic of aerial nontrophic and sporogenic hyphae^37, 39, 40, 43^. Visualization of colonies with fluorescently labeled HFBs under a fluorescent imaging system (ChemiDoc MP) revealed the complex HFB-determined architecture of the mycelia (Fig. 5). It was concordant with an abundant and coordinated intracellular accumulation of HFBs by aerial hyphae prior to sporulation. Then, the sporulation onsets were accompanied by the massive extracellular release of HFBs from nontrophic aerial hyphae and conidiophores. This process may explain how the colony efficiently prepares for conidiation because specific sites of mycelium where spores are to be formed become covered by a macroscopically visible HFB-rich protein matrix. We propose that it is produced to be rapidly and efficiently used for coating newly formed spores (*see below*).

In many molds, including *Trichoderma* spp., conidiophores are not evenly distributed on the colony surface but appear in pustules or form characteristic conidial rings in response to circadian rhythms^44,45^ or other stimuli^46^. We showed that in *Trichoderma* colonies, the emergence of such conidiation hot spots starts with the intracellular accumulation of HFBs (Supplementary material 6). We also observed high upregulation of multiple HFB-encoding genes at early stages of conidiogenesis (Fig. 5). Although only the *hfb4* and *hfb10* genes were involved in all stages of asexual development in both Th and Tg, the initiation of conidiation was linked to the short-term activity of at least two other *hfb* genes *(hfb2* and *hfb3).* The expression of all four genes decreased after the conidiophores were raised, and only *hfb4* and *hfb10* maintained their activity during spore formation, which is consistent with the particular importance of these two HFBs during sporulation. As reported previously^2^, mutants deprived of HFB4 and HFB10 produce significantly fewer spores than the parental strain; however, they still tend to form conidial rings or pustules. We assume that these proteins are required for normal (=abundant) conidiation, but the process itself is triggered by other factors, such as starvation, illumination or other stresses^44^.

Prior to their extracellular release, HFBs accumulate inside sporogenic hyphae. We propose that this accumulation is required for their normal development and the overall fitness of the colony^2^. The aerial hyphae of the *hfb*-deletion and *hfb*-overexpressing strains had signs of reduced and increased intracellular (turgor) pressure, respectively (Fig. 4). The apical hyphal tips of *hfb4*-deletion strains of Tg had characteristic indentations although these areas were known to have the highest turgor pressure^33^. This fact alone suggests the role of HFBs in turgor pressure. In contrast, HFB-overexpressing strains of both species and conidiophores of the wild-type strains had a visibly swollen appearance and were frequently covered by multiple large herniations filled with HFB-containing organelles or vesicles. Considering that the cell wall ultrastructure was not visibly changed, both observations indicated increased turgor pressure when HFB production was either naturally (conidiophore formation) or artificially (by using a strong constitutive promoter controlling *hfb* genes) stimulated. HFB-deletion strains were devoid of conidiation. We speculate that this phenotype was caused by an alteration of intracellular turgor pressure required for conidiophore emergence. To date, the understanding of turgor pressure in fungi is limited to studies of hydrophilic trophic hyphae frequently observed in fungi-like protozoans (Oomycota)^47^, plant-pathogenic fungi^48, 49^ or *Neurospora crassa*^50^ and explained by cytoplasm and water flow, osmosis and regulation of the mitogen-activated protein kinase cascade^33, 51^. However, these studies are not relevant to aerial hyphae where HFB-encoding genes are highly active. The mechanisms of turgor pressure in aerial hyphae are far not understood^52^. In aerial hyphae, turgor pressure is required not only for apical elongation but also for growth against the force of gravity and production of upregulated mechanically stable conidiophores. However, because aerial hyphae are usually hydrophobic (coated by HFBs or other surface-active proteins), their ability to absorb water and exchange ions within the environment is limited^52^ thus, the standard methods to measure turgor are not applicable. Because HFBs showed affinity to conidiophores and phialides, we propose that they contribute to the specific mechanisms for turgor pressure regulation in aerial hyphae and conidiophores. Detailed observations of conidiophore morphology in more than 200 *Trichoderma* spp. performed by W. M. Jaklitsch^53, 54^ showed that herniations of conidiophores are very common, at least in this genus. The accumulation of HFBs in tonoplasts, where they form or line up the membranes (*see below*), may promote their stability and could represent a major factor underlying turgor pressure formation in these specific cells. In this sense, the biological function of HFBs in *Trichoderma* observed in this study resembles the role of HFBs in the formation of mushrooms, where they provide hydrophobicity to fruiting bodies and contribute to the mechanical stability of aerial channels in basidiocarps^55, 56^. Interestingly, similar conclusions were proposed when the first hydrophobins in Ascomycota were discovered^15, 57^. Thus, we can generalize that intracellular HFBs are essential for the formation of sporogenic structures and therefore are universally present in all filamentous fungi regardless of the hydrophobicity of the hyphae and spores^12, 58^. Consequently, the accumulation and secretion of HFBs in the aerial mycelium largely explain the development and architecture of mold colonies.

Massive accumulation of highly surface-active molecules in a eukaryotic cell that has multiple compartments and membranes presents risks for normal metabolic processes. We revealed that HFBs tended to accumulate intracellularly not only in the aerial hyphae of a filamentous fungus but also in *P. pastoris* (regardless of the cultivation conditions, Fig. 2). Unexpectedly, despite the use of the secretory α-factor signal peptide from *S. cerevisiae*, the heterologous production of HFBs in *P. pastoris* resulted in intracellular accumulation, cell enlargement and release by cell bursting. Initially, we hypothesized that in both fungi, HFB-containing organelles filled up some of the cell compartments and influenced intracellular homeostasis. However, an expression analysis (Supplementary material 12) of marker genes of ER stress, autophagy, and the unconventional protein sorting pathway revealed that neither *P. pastoris* nor *Trichoderma* cells were stressed, which was also confirmed by the visually healthy phenotypes.

Due to the presence of the characteristic signal prepropeptide sequence, HFBs are initially translated into the ER lumen^26^, where the cysteine residues become oxidized and disulfide bonds are formed^59^. Our results suggest that in trophic and hydrophilic hyphae, where only a limited amount of HFB may be required and *hfb* genes are generally downregulated, HFBs are transported from the ER to vesicles and become conventionally secreted into the medium for lowering liquid surface tension. The accumulation of HFBs in aerial hyphae (where *hfb* genes are highly active) is probably started when these proteins become translocated from the ER to lipid bodies. The colocalization of fluorescently labeled HFBs and lipid droplets was observed in aerial hyphae and phialides of *Trichoderma* as well as in *P. pastoris* (Fig. 2). Previous studies have shown the ability of various HFBs to stabilize lipid emulsions^42, 60^. TEM analysis (Fig. 3) revealed that lipid droplets of HFB-overexpressing strains were more electron dense than those of the control strains, which may be caused by the enrichment of dissolved HFBs; however, this hypothesis requires further verification. Remarkably, Hähl et al.^32^ demonstrated that HFB1 from *T. reesei* (which is chemically similar to HFBs studied here^21^) can form pure bilayer HFB vesicles stable in a variety of liquids and sizes. Our observation that aerial hyphae and conidiophores in particular possessed large vacuoles or tonoplast-like structures that were massively HFB-positive indicated that intracellular HFBs formed similar membranous vesicles (HFB vesicles) as described in a cell-free system^32^. Comparisons of micrographs from superresolution fluorescent *in vivo* microscopy and TEM indicate that HFBs did not fill in these organelles but instead lined up their membranes. TEM micrographs of the wild-type, *hfb* deletion and HFB-overexpressing hyphae revealed multiple events of lipid body-to-vacuole internalization at an equal frequency in all strains, and such events were also reported for other fungi^31^. HFB-overexpressing strains were enriched in specific organelles that we assigned as VMSs putatively containing HFBs. Such vesicles were usually spherical and frequently included in one another, thus forming one large tonoplast resembling a “matryoshka”. We speculate that such structures were formed after internalization of HFB-enriched lipid droplets in vacuoles. In this case, HFBs likely assembled at the lipid/water interface and formed HFB membranes in VMSs. Importantly, the formation of VMSs with putative HFB vesicles was also observed in recombinant *P. pastoris* cells producing HFBs.

In *Trichoderma*, the release of HFB-enriched vesicles occurred first in the periplasm through the fusion (or budding) of VMSs with the plasma membrane. Consistently, the periplasm volume of *hfb*-deletion mutants was substantially reduced compared to that of the wild-type strains and HFB-overexpressing mutants. Our results do not allow us to conclude whether HFB membranes dissociate after/during the release from the cells; however, we observed extracellular HFB-enriched vesicles (Fig. 6 b). Previous reports stated that HFBs are secreted in a soluble form^7^. The solubility of HFB layers was used for grouping HFBs in conventional classes, where class I HFBs formed insoluble functional amyloids (rodlets), while the layers of class II HFBs (which were studied here) were soluble in organic solvents and ethanol^7, 11, 22^. However, Winandy et al.^42^ showed a similar stability of layers formed on glass by HFBs from both classes, and Hähl et al.^61^ demonstrated the stability of HFB vesicles formed by HFB1 (class II). Therefore, we speculate that HFBs, as universal and specific fungal proteins, contribute to the formation of extracellular vesicles that attract attention in cell biology as intracellular delivery and communication vehicles^62^, as demonstrated in fungi^63^, including *Trichoderma*^64^.

The subcellular vacuolar multicisternal structures similar to VMSs and fingerprint bodies detected in this study have been observed in many fungi, such as in studies characterizing the ultrastructure of *T. reesei* during cellulase production^65^ or investigating the benzyl-penicillin-overproducing strains of *Penicillium* sp.^66^. Their formation and the presence of membrane-covered lipid droplets were associated with a drastic increase in the cell wall thickness during penicillin biosynthesis. Structures resembling VMSs were also detected in the nematophagous fungus *Arthrobotrys oligospora*^67^ and in arbuscular mycorrhizal fungi^68^, and they are commonly seen in TEM micrographs^67^ of fungi producing HFBs from both classes.

The results of this study expand our understanding of the functions of vacuoles and lipid bodies in fungi and suggest that lipid internalization in vacuoles may offer a hydrophilic/hydrophobic interface for self-assembly of surface-active proteins, such as HFBs and possibly others (i.e., cerato-platanins^69^ or other SSCPs). Here, we showed that in specific cells, such as aerial hyphae, including conidiophores, vacuoles and lipid bodies serve as temporary storage for secreted proteins that require coordinated release and contribute to turgor pressure for arising fungal sporogenic structures. Moreover, the multiple herniations on the surface of conidiophores and aerial hyphae that were similar in size point to the heterogeneity and elasticity of the cell wall, and such herniations are common in *Trichoderma* spp.^54^, indicating rapid HFB secretion is possibly performed through squeezing out periplasmic HFB vesicles through the cell wall.

The superior surface activity and lack of toxicity^7^ make HFBs attractive proteins for various biomedical and biotechnological applications^39, 41^. Our findings of massive secretion of class II HFBS by aerial hyphae and intracellular retention of these proteins in fungal cells offer insight for effective production protocols. Even more interestingly, the verification of our hypothesis that stable HFB vesicles occur in living systems may be explored in synthetic biology^32^, where such vehicles are highly desired for drug delivery.

## Materials and Methods

### Strains and cultivation conditions

All strains used in this study are listed in Supplementary material 2. If not otherwise specified, filamentous fungi were maintained on PDA (Sigma, USA) at 25 °C in dark conditions. *Komagataella pastoris* (Saccharomycetales, syn. *Pichia pastoris)* was maintained on yeast extract peptone dextrose medium (YPD) at 28 °C. Yeasts were cultivated in buffered minimal glycerol medium (BMG, including 100 mM potassium phosphate, pH 6.0, 1.34% yeast nitrogen base, 4 × 10^−5^% biotin and 1% glycerol) and then transferred to buffered minimal methanol medium (BMM, including 100 mM potassium phosphate, pH 6.0, 1.34% yeast nitrogen base, 4 × 10^−5^% biotin and 0.5% methanol) for the designed protein production. All strains are available from the TU Collection of Industrial Microorganisms (TUCIM) at TU Wien, Vienna (Austria) and Nanjing Agricultural University, Nanjing (China).

### Molecular biology techniques

#### Gene expression

For the gene expression analysis, fungal biomass corresponding to the four stages of the life cycle, namely, (i) germlings, (ii) trophic hyphae, (iii) aerial hyphae, and (iv) conidiophores and conidia, were collected from cellophane-covered agar plates or liquid cultures. Total RNA was extracted using an RNeasy Plant MiniKit (Qiagen, Germany) according the manufacturer’s protocol. cDNA was synthesized with a RevertAid™ First Strand cDNA Kit (Thermo Fischer Scientific, USA) using the oligo (dT)^18^ primer. qPCR was performed using qTOWER (Jena Analytics, Germany) for the genes of interest (listed in Supplementary material 1) and calculated by the 2^ΔΔCt^ method^70^ using *tef1* as the housekeeping gene^71^. PCRs were prepared in a total volume of 20 μl containing 10 μl of iQ SYBR Green Supermix (Bio-Rad, USA), 0.5 μM each primer and 100 ng of cDNA. The program was 40 cycles with 30 s at 95 °C and 60 s at 60 °C, which was initially denatured for 6 min at 95 °C. A melting curve from 55 °C to 95 °C was generated for each qPCR run.

#### Gene deletion and reverse complementation

The polyethylene glycol (PEG)-mediated protoplast transformation procedure was adopted^71^. Gene deletion in Tg and Th was performed as described previously in Cai et al.^2^ using a hygromycin B cassette (*hph*, from the plasmid pPcdna1-hph^72^) and/or geneticin cassette *(neo*, from the plasmid pPki-Gen^73^) to replace the gene of interest (Supplementary material 17). For reverse complementation, a copy of the gene with its promoter and terminator region from the wild-type strain or a copy of this gene with a fluorescent tag *(mrfp* or *yfp*, see below) fused at the C-terminus was amplified and inserted into the corresponding deletion mutant. The positive mutants were further confirmed at the transcriptional level. All primers used in this study are given in Supplementary material 18.

#### *In vivo* fluorescent labeling

HFB-encoding genes were conjugated with genes encoding fluorescent proteins (followed by a 6×His tag) at the C-terminus over a synthetic GGGGS×3 linker^74^. Briefly, a 1.2 kb fragment including the entire *hfb* gene before the stop codon (namely, the 5’ homologous arm), a 1.0 kb fragment of its native terminator region and a 1.2 kb fragment after the terminator region (namely, the 3’ homologous arm) were amplified by PCR and then cloned into the pUC19 plasmid (Thermo Fisher Scientific, USA) in the order of the 5’ homologous arm, the fluorescent protein gene (either yellow fluorescent protein (YFP; excitation at 514 nm and emission at 527 nm) or mutated red fluorescent protein (mRFP; excitation at 559 nm and emission at 630 nm), terminator region, selection marker cassette and 3’ homologous arm by a ClonExpress® MultiS One Step Cloning Kit (Vazyme, China). The obtained plasmids were confirmed by Sanger sequencing to determine the in-frame fusion constructs and transformed into *Trichoderma* with the above two selection markers. For the putatively labeled strains, transformants from PEG-mediated protoplast transformation were screened by PCR using the same strategy as above and confirmed by the presence of the corresponding fluorescent signals by a Leica DMi8 microscope with fluorescent applications (Leica, Germany). Additionally, a mutant expressing mRFP under the control of an *hfb* promoter (P_*hfh4*_) was generated. All PCR products and vector constructs were verified by sequencing. Details on the mutant construction process are provided in Supplementary material 17.

#### Overexpression of *hfb* genes in *Trichoderma*

Overexpression vectors were constructed by cloning a 1.6 kb fragment that contained the open reading frame (ORF) of the *hfb* gene and its terminator region into the *pUCPcdna1-hph* plasmid after a constitutive promoter, *P_cdna1_*, from *T. reesei* QM 6a^72^. The PCR products were purified and fused into a ClaI-digested plasmid with an In-Fusion HD cloning kit (TAKARA, Japan). Mutants for overexpressing HFB2 (_Tg_OPB38530) fused with a mRFP tag were generated with the fluorescent construct mentioned above.

#### Heterologous expression of *hfb* genes from *Trichoderma* in *P. pastoris*

An EasySelect™ *Pichia* Expression Kit was used to express genes from *Trichoderma* in the *P. pastoris* strain KM71H (Invitrogen, USA) according to the manufacturer’s instructions. The encoding region after the end of the predicted signal peptide of the *hfb* gene (SignalP 4.1) to the stop codon was amplified from the cDNA of the wild-type strain. The cloned fragment was inserted into the plasmid pPICZαA containing the AOX1 promoter by replacing the fragment between the EcoR I site and Xba I site using an In-Fusion HD cloning kit (Clontech, Japan). To express recombinant proteins with a fluorescent tag, a fluorescent protein-encoding gene (a green fluorescent protein variant (i.e., GFPuv, excitation at 395 nm and emission at 509 nm) was inserted at the C-terminus of the designated *hfb* gene. Additionally, the recombinant protein encoded by this construct contained the *Saccharomyces cerevisiae* α-mating factor prepropeptide^75^ at the N-terminus of the *hfb* gene and a His×6 epitope at the C-terminus. The resulting vector was linearized with Sac I or Pme I and then transformed into *P. pastoris* by electroporation. One of the immunoblotting-positive transformants (see below) was chosen to obtain the recombinant protein.

#### Biochemical techniques

For sodium dodecyl sulfate-polyacrylamide gel electrophoresis (SDS-PAGE) and immunoblotting assays, protein samples from the surface of *Trichoderma* spores or hyphae were collected by washing with 1% SDS (pH 8.1, 50 mM HCl-Tris). The *Pichia*-produced proteins were collected from BMM fermentation (see *P. pastoris* fermentation in Przylucka et al.^39^). Protein samples were then denatured and loaded into a 15% polyacrylamide gel, followed by silver staining with a SilverQuest™ Silver Staining Kit (Life Technologies, Germany) suitable for HFBs^39^. Immunological visualization of HFBs was performed using a mouse anti-His-tag horseradish peroxidase (HRP) antibody (Genescript, USA). The target proteins were visualized following the protocol supplied with Clarity™ Western ECL (Bio-Rad, USA) or with the ONE-HOUR Western™ Standard Kit (Genscript, China).

### Phenotype investigation

#### Macromorphology

The strains were cultivated on PDA plates for seven days at 25 °C in darkness. The colony morphology was recorded by a Cannon EOS 70D (equipped with a Cannon 100 μm macro lens) under white light. The development of the colonies of the fluorescently labeled *Trichoderma* strains was monitored by making regular (every 24 h) images in a Bio-Rad ChemiDoc MP system (Bio-Rad, USA) equipped with multiplex fluorescent channels. The system was first optimized based on the florescence of single labeled strains and then applied for the double labeled mutants.

#### *In vivo* microscopy and fluorescent staining

The fluorescently labeled strains were imaged by a stereo confocal microscope (Olympus, MVX10, Japan). Protein localization in fungal cells was carried out by an UltraVIEW VoX Spinning Disk Confocal Microscope (PerkinElmer, USA) and a Leica DMi8 (Leica, Germany), and 3D images and videos were acquired by a Leica LAS X (Germany). The detailed intracellular localization of HFBs was performed by a Fast Super-Resolution Laser Confocal Microscope (Zeiss LSM980 Airyscan2, Germany). For intracellular lipid staining, fungal cells were either incubated in 2 μM BODIPY™ FL C_12_ (excitation 500 nm, emission 510 nm, Thermo Fischer Scientific, USA) for 15 min and washed with 50 mM PBS three times or incubated in 1× HCS LipidTOX Green Phospholipidosis Detection Reagent (excitation 495 nm, emission 525 nm, Thermo Fischer Scientific, USA) for 5 min. Stained samples were imaged using the Leica DMi8 microscope (Leica, Germany) or the Fast Super-Resolution Laser Confocal Microscope.

#### Electron microscopy

Fungal colonies were investigated by cryo-SEM (Quorum PP3010T integrated onto a Hitachi SU8010 FE-SEM, Japan). The culture samples were rapidly frozen in nitrogen slush, fractured at −140 °C and coated with 5 nm platinum.

The cell ultrastructure was investigated by TEM (Hitachi H-7650, Japan). Fungal cells were fixed in a 2.5% glutaraldehyde solution at 4 °C overnight. Fixed cells were then washed with 0.1 M PBS three times, postfixed with 1% osmium tetroxide (for 2 h), washed with 0.1 M PBS again and dehydrated with a gradation of ethanol, namely, 50%, 70%, 90% and 100%, followed by 100% acetone. Dehydrated samples were infiltrated in graded acetone/epoxy resin and cured at 60 °C for 48 h. Cured resin blocks were trimmed, sectioned at a 70-nm thickness and poststained with uranyl acetate and lead citrate (3%).

#### Statistical analyses

The data were calculated and statistically examined by one-way analysis of variance (ANOVA) or multivariate analysis of variance (MANOVA) using STATISTICA 6 (StatSoft, Germany). Heatmap analysis and average linkage hierarchical clustering, and PCA plots were obtained using R (version 3.2.2). The significance level was set at *p* < 0.05 unless otherwise stated.

## Supporting information

Supplementary materials 1-9, 11-14, 17. Supplementary results and detailed methods.

Supplementary material 10. Intracellular localization of HFB2::mRFP in two-day-old aerial hyphae of the TgOEhfb2::mrfp strain.

Supplementary material 15. Animated 3D reconstructions of extracellular HFB-enriched matrices coating sporulating Trichoderma colonies.

Supplementary material 16. Video showing putative HFB-enriched vesicles secreted by the TgOEhfb2::mrfp strain.

Supplementary material 18. Primers used in this study.

## Data availability

All relevant data are available from the authors. The source data underlying Fig. 5, and Supplementary material 1, 9, 12 and 17 are provided as a Source Data file.

## Acknowledgments

The research in China was supported by grants from the National Natural Science Foundation of China (KJQN201920) and the Ministry of Science & Technology of Jiangsu Province (BK20180533) to FC. The research in Austria was supported by the Austrian Science Foundation (FWF), P25613-B20, to ISD and the Vienna Science and Technology Fund (WWTF), LS13-048, to ISD. The numerical calculations of MD simulations reported in this paper were partially performed at TUBITAK ULAKBIM, High Performance and Grid Computing Center (TRUBA resources) and Acibadem Mehmet Ali Aydinlar University High Performance Computing Server.

The authors are thankful to Dr. Lihui Wei (Jiangsu Academy of Agricultural Science, Nanjing, China) for the use of a stereo confocal microscope. We thank Yan Teng at the Institute of Biophysics, CAS, for providing technical support for superresolution CLSM. We are very grateful to Christian P. Kubicek (Vienna, Austria) for performing a critical reading of the manuscript drafts and engaging in discussions.

## Authors’ contributions

FC carried out experiments with the assistance of ZZ, RG, MD, SJ, KC, QG, JZ, AP, GBA and ISD. PX and ZF provided expertise on superresolution CLSM. GBA performed the *in silico* analysis of fluorescently labeled HFBs. ISD and QS conceived and designed the study. ISD and FC carried out the data analysis, prepared the figures and supplements, and wrote the manuscript with comments from QS and GBA. All authors read and approved the manuscript.

## Supplementary materials

**Supplementary materials 1-9, 11-14, 17.** Supplementary results and detailed methods.

**Supplementary material 10.** Intracellular localization of HFB2::mRFP in two-day-old aerial hyphae of the _Tg_*OEhfb2::mrfp* strain.

**Supplementary material 15.** Animated 3D reconstructions of extracellular HFB-enriched matrices coating sporulating *Trichoderma* colonies.

**Supplementary material 16.** Video showing putative HFB-enriched vesicles secreted by the _Tg_*OEhfb2::mrfp* strain.

**Supplementary material 18.** Primers used in this study.

