## Supplementary materials 1-9, 11-14, 17. Supplementary results and detailed methods. for "Intracellular accumulation and secretion of hydrophobin-enriched vesicles aid the rapid sporulation of molds"

**Cai et al.**

**Running title: Intracellular functions of hydrophobins in molds**

Feng Cai^1-3^, Zheng Zhao^2^, Renwei Gao^2^, Mingyue Ding^2^, Siqi Jiang^2^, Qi Gao^1^, Komal Chenthamara^3^, Marica Grujic^3^, Zhifei Fu^4^, Jian Zhang^1^, Agnes Przylucka^3^, Pingyong Xu ^4^, Günseli Bayram Akcapinar^3,5^, Qirong Shen^1*^, Irina S. Druzhinina^1-3*^

^1^The Key Laboratory of Plant Immunity, Jiangsu Provincial Key Lab of Solid Organic Waste Utilization, Nanjing Agricultural University, Nanjing, China

^2^Fungal Genomics Laboratory (FungiG), Nanjing Agricultural University, Nanjing, China

^3^Institute of Chemical, Environmental and Bioscience Engineering (ICEBE), TU Wien, Vienna, Austria

^4^Key Laboratory of RNA Biology, Institute of Biophysics, Chinese Academy of Science, Beijing, China

^5^Department of Medical Biotechnology, Institute of Health Sciences, Acibadem Mehmet Ali Aydinlar University, Istanbul, Turkey

Content

### Supplementary material 1 *hfb* expression pattern during *Trichoderma* development

**Table S1.1** Expression of *hfb*-encoding genes during four stages of *Trichoderma* development

| **Parental strain   NCBI Genome ID** | ***hfb*  gene** | **NCBI Accession Numbers** | | **Relative gene expression** | | | | | | | |
| --- | --- | --- | --- | --- | --- | --- | --- | --- | --- | --- | --- |
|  |  |  |  | **Spore germination** | | **Trophic hyphae** | | **Aerial hyphae** | | **Conidiation** | |
|  |  |  |  | **Fold  expression** | SD | **Fold  expression** | SD | **Fold  expression** | SD | **Fold  expression** | SD |
| ***T. harzianum* CBS 226.95  (MBGI00000000.1)** | n.a.* | PTB60449 | MF527127 | **0** |  | **0** |  | **0** |  | **0** |  |
|  | *hfb3* | PTB52129 | MF527128 | **0** |  | **7.02E-05** | *2.9E-05* | **4.93E-01** | *1.6E-01* | **6.14E-02** | *7.4E-03* |
|  | ***hfb4*** | PTB58174 | MF527129 | **4.67E-05** | *9.0E-06* | **3.11E-04** | *8.9E-06* | **3.47E+00** | *3.7E-02* | **1.27E+00** | *2.2E-01* |
|  | *hfb5* | PTB53925 | MF527130 | **0** |  | **0** |  | **0** |  | **0** |  |
|  | *hfb6* | PTB60167 | MF527131 | **0** |  | **0** |  | **0** |  | **0** |  |
|  | ***hfb2***** | PTB60601 | MF527132 | **0** |  | **1.08E-01** | *1.9E-02* | **1.06E+00** | *2.1E-01* | **1.75E-01** | *2.3E-02* |
|  | *hfb9a* | PTB50855 | MF527133 | **0** |  | **9.43E-06** | *4.3E-06* | **2.37E-05** | *6.7E-06* | **1.73E-03** | *3.6E-04* |
|  | *hfb9b* | PTB48391 | MF527134 | **0** |  | **6.62E-06** | *1.8E-06* | **2.26E-04** | *1.3E-04* | **5.95E-04** | *6.1E-05* |
|  | ***hfb10*** | PTB48206 | MF527135 | **2.85E-01** | *2.5E-02* | **1.47E-01** | *4.6E-02* | **2.14E+00** | *3.0E-01* | **7.66E-01** | *1.1E-01* |
|  | n.a. | PTB49111 | MF527136 | **0** |  | **8.54E-04** | *7.9E-05* | **5.23E-04** | *1.1E-04* | **5.39E-05** | *1.4E-05* |
|  | n.a. | PTB56946 | MF527138 | **0** |  | **0** |  | **0** |  | **0** |  |
| ***T. guizhouense* NJAU 4742  (LVVK00000000.1)** | *hfb3* | OPB45549 | MF527117 | **0** |  | **9.68E-03** | *1.1E-03* | **3.19E-01** | *4.5E-02* | **1.59E-02** | *3.8E-03* |
|  | ***hfb4*** | OPB37525 | MF527118 | **1.58E-04** | *6.2E-05* | **1.12E-02** | *2.5E-03* | **1.30E+00** | *1.6E-01* | **7.36E-01** | *1.7E-01* |
|  | *hfb5* | n.a. | MF527119 | **0** |  | **1.22E-04** | *4.1E-05* | **4.88E-03** | *5.8E-04* | **3.18E-02** | *5.2E-03* |
|  | *hfb6* | OPB38878 | MF527120 | **0** |  | **0** |  | **0** |  | **0** |  |
|  | ***hfb2***** | OPB38530 | MF527121 | **0** |  | **1.87E-01** | *1.5E-02* | **7.66E-01** | *1.2E-01* | **2.83E-02** | *1.0E-02* |
|  | *hfb9a* | OPB40515 | MF527122 | **0** |  | **6.32E-06** | *1.4E-06* | **6.58E-06** | *3.6E-06* | **3.69E-04** | *9.0E-05* |
|  | *hfb9b* | OPB44528 | MF527123 | **0** |  | **3.39E-03** | *6.9E-04* | **7.15E-03** | *1.1E-03* | **2.35E-03** | *3.7E-04* |
|  | ***hfb10*** | OPB44696 | MF527124 | **1.90E+00** | *8.0E-01* | **1.31E+01** | *3.7E-01* | **1.42E+01** | *2.7E+00* | **2.34E+00** | *2.2E-01* |
|  | n.a.* | OPB45278 | MF527125 | **0** |  | **3.53E-06** | *2.2E-06* | **1.73E-05** | *5.9E-06* | **0.0019** | *2.6E-04* |

*n.a., not available

***hfb2*, representing *hfb2* sensu lato including *hfb7^1, 2^*.

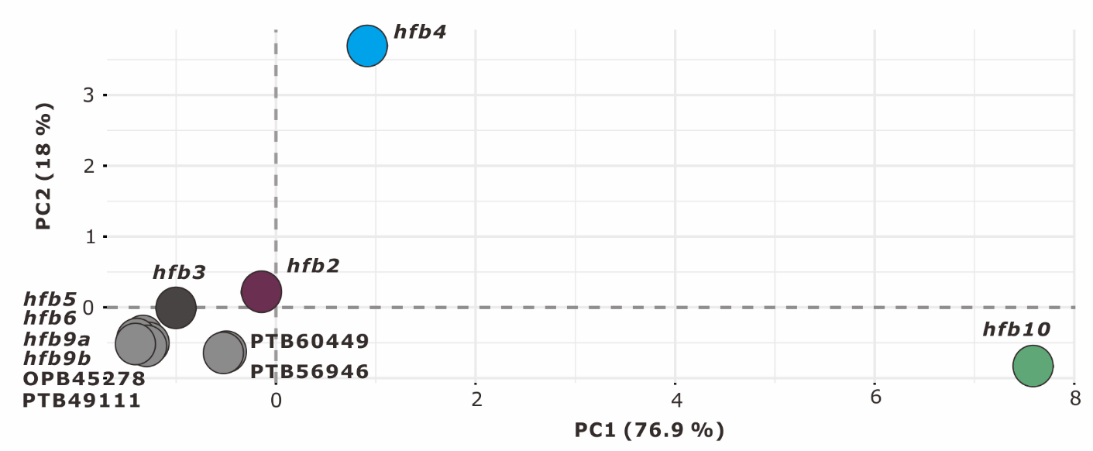

**Figure S1.1** Principal component analysis (PCA) of the *hfb* expression pattern in *T. harzianum* and *T. guizhouense*. The complete list of *hfb*-encoding genes present in both genomes is given in Table S1.

### Supplementary material 2 Strains used in this study

**Table S2.1** Strains used in this study

| **Strain description** | **TUCIM ID** | **Strain name** | **TUCIM ID** | **Strain name** |
| --- | --- | --- | --- | --- |
|  | ***Trichoderma guizhouense*** | | ***Trichoderma harzianum*** | |
| Wild type | 4742 | **NJAU 4742** | 916 | **CBS 226.95** |
| Fluorescently labeled mutants | 7436 | **_Tg_*hfb4*::*mrfp*-*hfb10*::*yfp*** | 7422 | **_Th_*hfb4*::*yfp-hfb10*::*mrfp*** |
| *hfb4*-deleted mutant | 7435 | **_Tg_Δ*hfb4*-1** | 7421 | **_Th_Δ*hfb4*-3** |
|  | 7434 | **_Tg_Δ*hfb4*-4** | 7420 | **_Th_Δ*hfb4*-11** |
| *hfb10*-deleted mutant | 7431 | **_Tg_Δ*hfb10*-2** | 7417 | **_Th_Δ*hfb10*-2** |
|  | 7430 | **_Tg_Δ*hfb10*-3** | 7416 | **_Th_Δ*hfb10*-17** |
| *hfb4* and *hfb10* double-deleted mutant | 7426 | **_Tg_Δ*hfb4*-Δ*hfb10*-2** | 7413 | **_Th_Δ*hfb4*-Δ*hfb10*-27** |
|  | 7427 | **_Tg_Δ*hfb4*-Δ*hfb*10-11** | 7412 | **_Th_Δ*hfb4*-Δ*hfb10*-30** |
| *hfb2*-deleted mutant | 7442 | **_Tg_Δ*hfb2*-1** | 8005 | **_Th_Δ*hfb2*-1** |
|  | 7441 | **_Tg_Δ*hfb2*-17** | 8006 | **_Th_Δ*hfb2*-2** |
| *hfb4*-overexpressing mutant | 7433 | **_Tg_OE*hfb4*-6** | 7419 | **_Th_OE*hfb4*-11** |
|  | 7432 | **_Tg_OE*hfb4*-13** | 7418 | **_Th_OE*hfb4*-13** |
| *hfb10*-overexpressing mutant | 7429 | **_Tg_OE*hfb10*-9** | 7415 | **_Th_OE*hfb10*-1** |
|  | 7428 | **_Tg_OE*hfb10*-10** | 7414 | **_Th_OE*hfb10*-6** |
| *mrfp*-labeled *hfb2*-overexpressing mutant | 7439 | **_Tg_OE*hfb2*::*mrfp*-23** | n.a. | |
|  | 7438 | **_Tg_OE*hfb2*::*mrfp*-24** |  |  |
| *mrfp*-labeled *hfb3* strain | 10530 | **_Tg_*hfb3*::*mrfp*** |  |  |
| *mrfp*-labeled *hfb4* strain | 7424 | **_Tg_*hfb4*::*mrfp*** |  |  |
| *ypt7*-deleted mutant | 8001 | **_Tg_Δ*ypt7*-30** |  |  |
|  | 8002 | **_Tg_Δ*ypt7*-55** |  |  |
| *mrfp*-labeled *hfb4-* and *ypt7-*deleted mutant | 8003 | **_Tg_*hfb4*::*mrfp*-Δ*ypt7*-11** |  |  |
|  | 8004 | **_Tg_*hfb4*::*mrfp*-Δ*ypt7*-17** |  |  |
| *hfb4*-complementary mutant to *hfb4-*deleted mutant | 10522  10523 | **_Tg_Δ*hfb4hfb4*-10**  **_Tg_Δ*hfb4-hfb4*-15** |  |  |
| *hfb4*::*mrfp*-complementary mutant to *hfb4-*deleted mutant | 10524  10525 | **_Tg_Δ*hfb4*::*hfb4*::*mrfp*-3**  **_Tg_Δ*hfb4*::*hfb4*::*mrfp* -5** |  |  |
| *hfb10*-complementary mutant to *hfb10-*deleted mutant | 10526  10527 | **_Tg_Δ*hfb10*::*hfb10*-11**  **_Tg_Δ*hfb10*::*hfb10-*12** |  |  |
| *hfb10*::*ypf*-complementary mutant to *hfb10-*deleted mutant | 10528  10529 | **_Tg_Δ*hfb10*::*hfb10*::*yfp*-1**  **_Tg_Δ*hfb10*::*hfb10*::*yfp*-7** |  |  |
| *mrfp-*expressing strain under the control of the *hfb4* promoter | 8000 | **_Tg_P*_hfb4_*::*mrfp*** |  |  |
| ***Pichia pastoris*** | | | | |
| Wild type | | | 8007 | **KM71H** |
| KM71H strain transformed with the empty vector of pPICZαA | | | 6633 | **_Pp_vec** |
| *hfb4-*expressing strain under control of the *P*_aox1_ promoter with the signal peptide of α mating factor from *Saccharomyces cerevisiae* | | | 6616 | **_Pp_P*_aox1_S*_α_::*hfb4*** |
| *hfb2-*expressing strain under control of the *P*_aox1_ promoter with the signal peptide of α mating factor from *S. cerevisiae* | | | 6617 | **_Pp_P*_aox1_*S_α_*::hfb2*** |
| *gfpuv-*labeled *hfb4-*expressing strain under control of the *P*_aox1_ promoter with the signal peptide of α mating factor from *S. cerevisiae* | | | 6625 | ***_Pp_P_aox1_S_α_*::*hfb4*::*gfpuv*** |
| *mrfp-*labeled *hfb2-*expressing strain under control of the *P*_aox1_ promoter with the signal peptide of α mating factor from *S. cerevisiae* | | | 6626 | **_Pp_P*_aox1_*S_α_::*hfb2*::*mrfp*** |

n.a., not available.

TUCIM, the TU Collection of Industrial Microorganisms, Vienna, Austria.

### Supplementary material 3 Molecular dynamic simulation and solvent accessible surface area calculation of fluorescently tagged HFBs used in this study

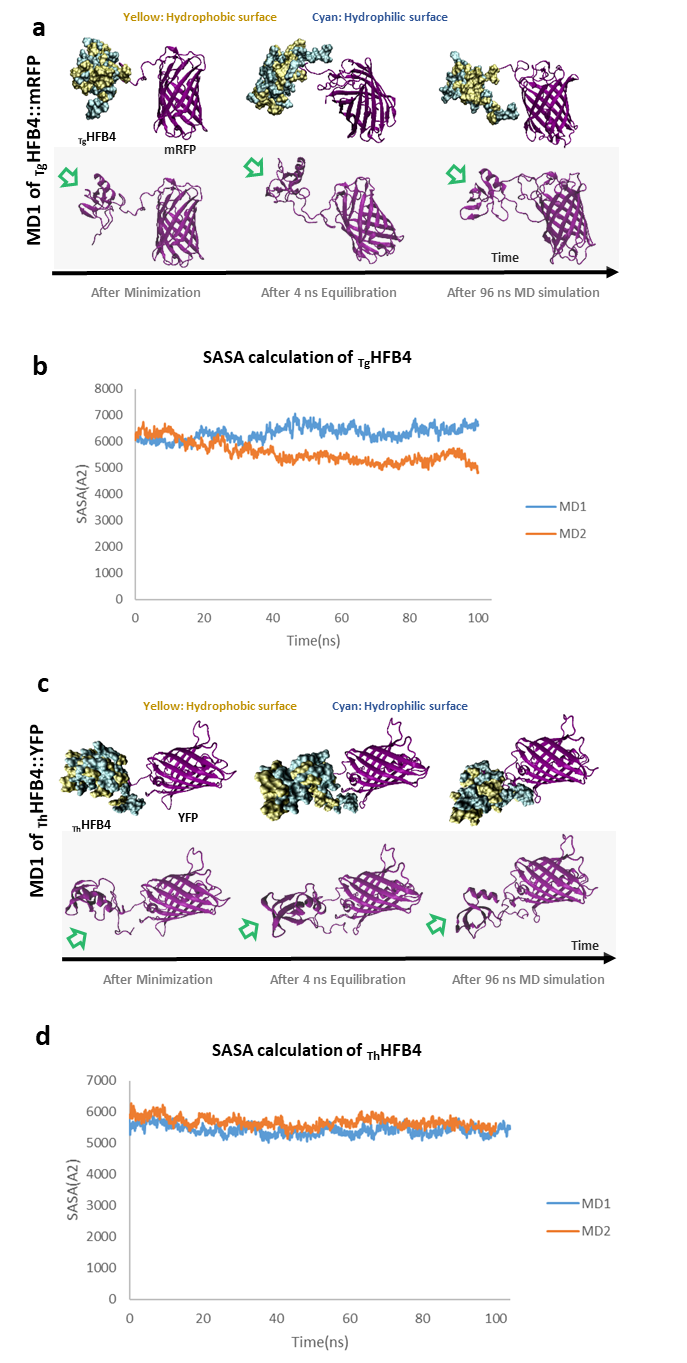

**Figure S3.1** Molecular dynamic (MD, **a** and **c**) simulation and solvent accessible surface area (SASA, **b** and **d**) calculation showed no putative disrupting effect of the fluorescent tags on the studied HFB4. MD simulations were used to analyze the effects of the fusion partners (mRFP and YFP) on the HFB4 protein structures (_Tg_HFB4 and _Th_HFB4), respectively, by the NAMD/VMD software package^3^. Simulations were performed inside a 10 Å water box under periodic boundary conditions at 298 °K using TIP3P water. All the structures were neutralized by the addition of Na^+^ or Cl^-^ ions. 2 fs timestep was used and data collection was done every 2 ps. Modelled structures were minimized 50000 steps using conjugate gradient (CG) method before the simulations. 100 ns (4 ns equilibration and 96 ns production runs) MD simulations were performed at 298 °K using the NPT ensemble under constant pressure and temperature. Root mean square deviation (RMSD) is adopted to indicate the large structural changes in the protein and to measure the scalar distance between atoms of the same type for two structures^4^. SASA is used to calculate the surface area of an atom, a residue, a molecule which is exposed to a specific solvent. This factor is measured in angstroms2 (Å2)^4^. SASA of the two fusion proteins (_Tg_HFB4::mRFP and _Th_HFB4:: YFP) were calculated for each fusion partner along the simulation time from the trajectories of the MD simulations performed at 298 K using water sphere with a radius of 1.4 Å via VMD scripting. Homology modelling of was performed using Modeller9v23^5, 6^ based on the HFBII structure deposited in protein databank (PDB ID: 1R2M-Chain A) from *T. reesei* for _Tg_HFB4 and _Th_HFB4, and mCherry structure (PDB ID: 6B0B-Chain D) and GFP structure (PDB ID: 4Xl5-Chain A) for mRFP and YFP, respectively. Ten different models were generated for each fusion protein and compared. Best scoring models are selected for the subsequent MD simulation. Note: MD were performed with two repeat runs (MD1 and MD2) and here only MD1 is shown as a representative example. Green arrows point to the hydrophobic patch of HFBs.

### Supplementary material 4 Response of *hfb* regulation in *Trichoderma* to the *hfb* gene manipulation

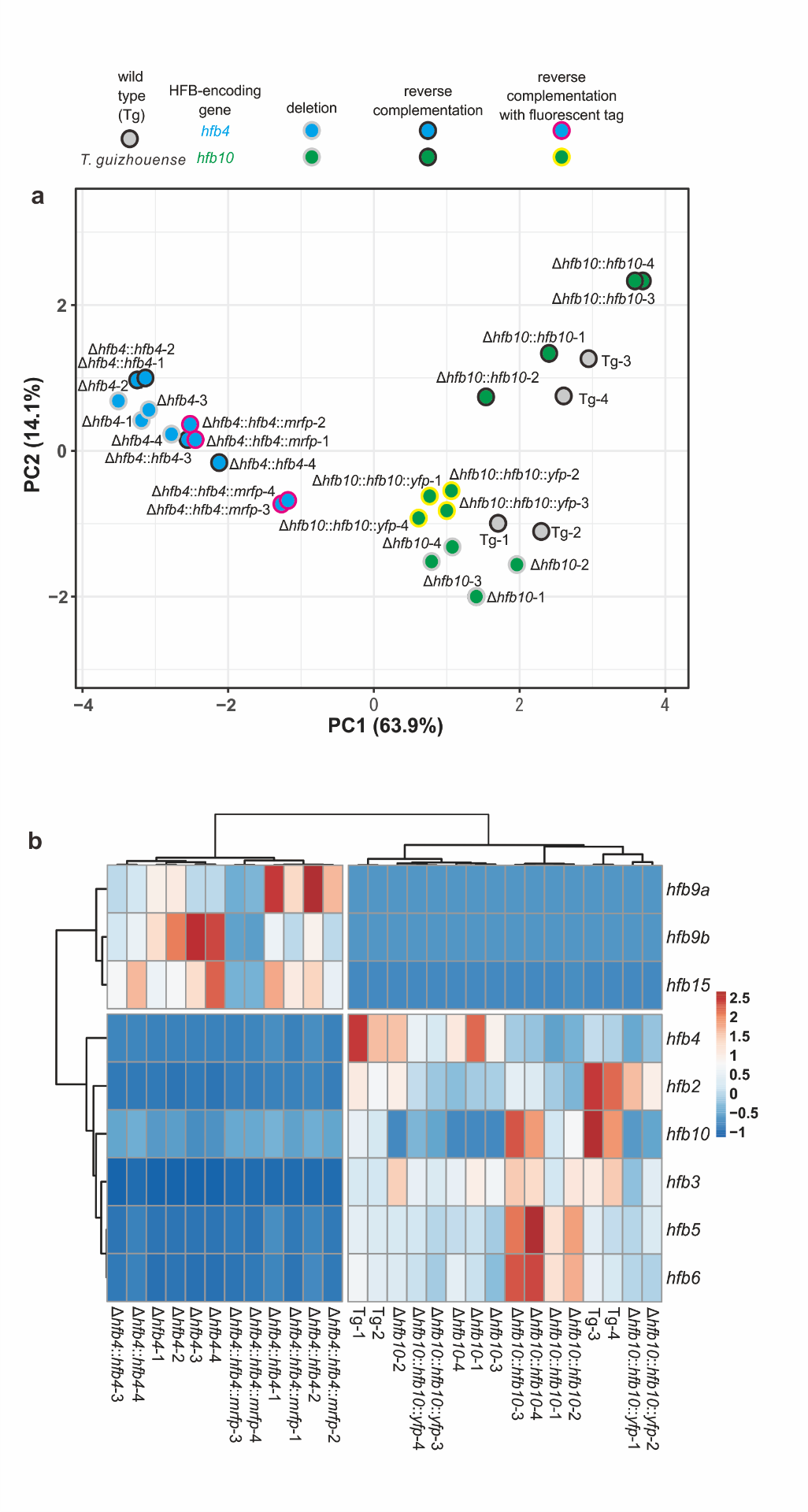

**Figure S4.1** Expression pattern of HFB-encoding genes in *T. guizhouense* NJAU 4742 (Tg) and its *hfb4*- or *hfb10*-related mutants. **a**, Principal component analysis (PCA) of the *hfb* expression pattern in all tested strains*.* Blue circles represent *hfb4*-related mutants, including *hfb4*-deletion (highlighted with gray periphery, Δ*hfb4*) and *hfb4*-deletion mutants complemented with *hfb4* from Tg (highlighted with black periphery, Δ*hfb4*::*hfb4*) or its labeled construction *hfb4*::*mrfp* (highlighted with red periphery, Δ*hfb4*::*hfb4::mrfp*). Green circles represent *hfb10*-related mutants, including *hfb10*-deletion (highlighted with gray periphery, Δ*hfb10*) and *hfb10*-deletion mutants complemented with *hfb10* from Tg (highlighted with black periphery, Δ*hfb10*::*hfb10*) or its labelled construction *hfb10*::*yfp* (highlighted with yellow periphery, Δ*hfb10*::*hfb10*::*yfp*). **b**, heatmap of HFB-encoding gene expression pattern of strains presented in **a**. The reverse complemented mutants where tagged and untagged HFB-encoding genes showed that the tag of mRFP and YFP had insignificant impact on the expression pattern of *hfb* genes. Importantly, although the re-introduction of the original copy of the gene (*hfb4* or *hfb10*) to the respective deletion mutant resulted into a comparable transcription of the respective gene relative to that of the wt, the expression pattern of the other *hfb* members was not recovered (as in the wt), indicating an interactive regulation network of *hfb* gene family .

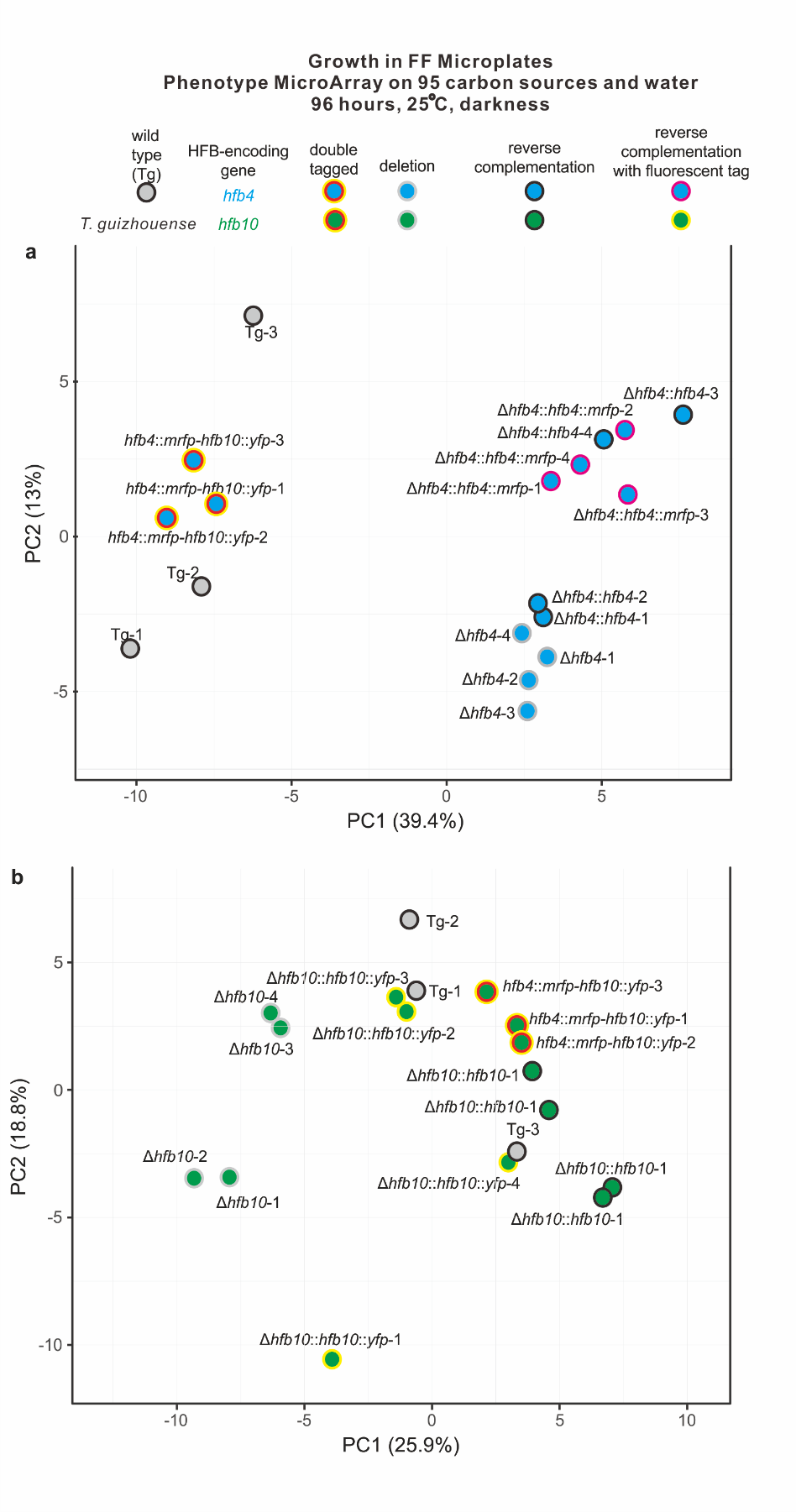

**Figure S4.2** Phenotypes of HFB-encoding genes in *T. guizhouense* NJAU 4742 (Tg) and its *hfb4*-(**a**) or *hfb10* (**b**)-related mutants estimated using the principal component analysis (PCA) of the Biolog Phenotype Microarrays data collected after 96 hours of incubation at 25°C in darkness as described in Cai et al.^10^*.* Symbols correspond to those in **Figure S4.1**. In addition, the double labeled strains are shown with red and yellow peripheries. The reverse complemented mutants where tagged and untagged HFB-encoding genes showed that the tag of mRFP and YFP had insignificant impact on the phenotypes of the respective strains. Importantly, the re-introduction of the original copy of the gene (*hfb4* or *hfb10*) to the respective deletion mutant resulted into a comparable growth profile of the respective untagged mutant relative to that of the wt.

### Supplementary material 5 Phenotypic characterization of mutants

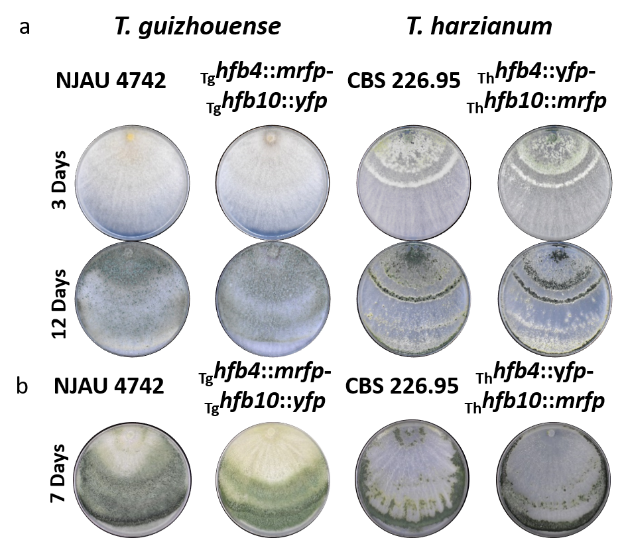

**Figure S5.1** Representative morphology of the wild type strains of Trichoderma (T. guizhouense NJAU 4742 and T. harzianum CBS 226.95) and their respective hfb4 and hfb10 double labelled mutants used in this study. Trichoderma strains grown on PDA at 25 °C.

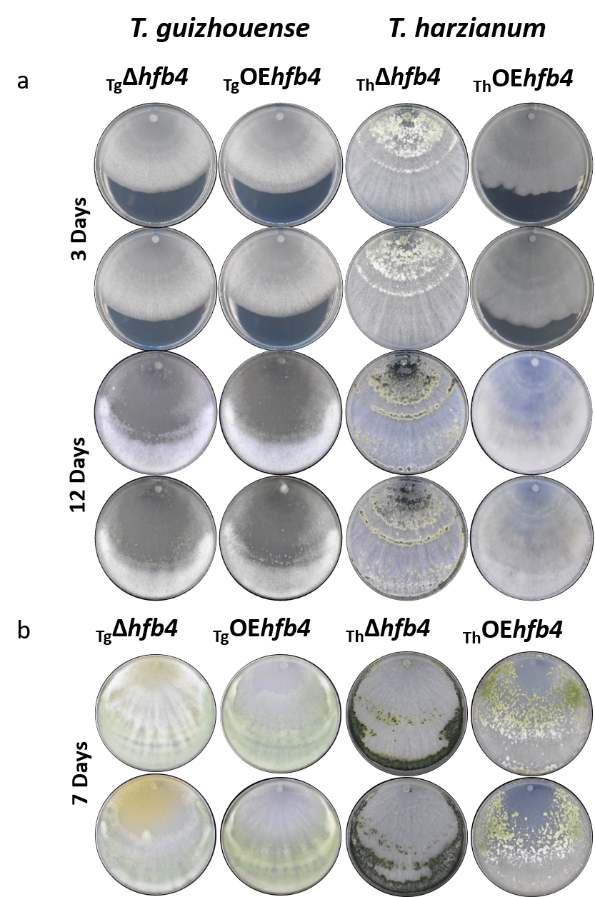

**Figure S5.2** Representative morphology of Trichoderma hfb4 mutants (N=2 for each genotype) generated in this study. Trichoderma strains grown on PDA at 25 °C.

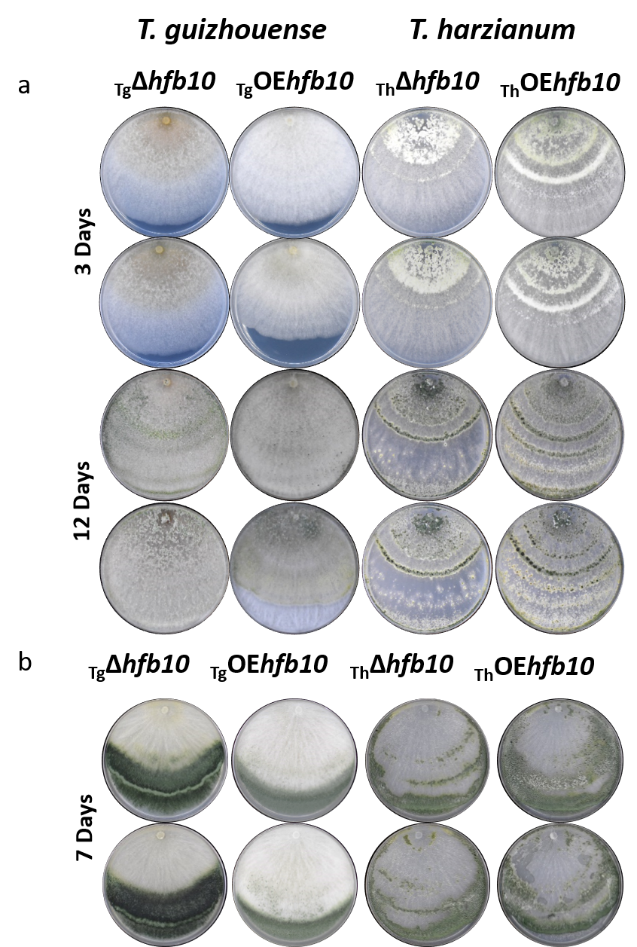

**Figure S5.3** Representative morphology of Trichoderma hfb10 mutants (N=2 for each genotype) generated in this study. Trichoderma strains grown on PDA at 25 °C.

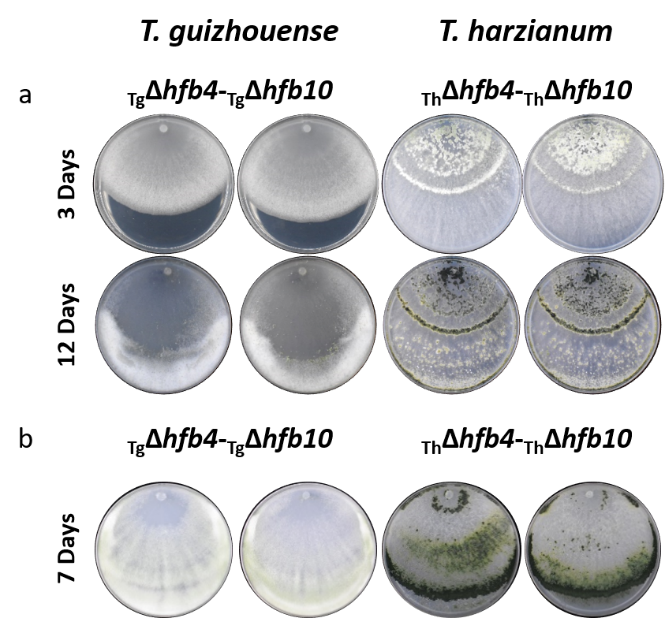

**Figure S5.4** Representative morphology of Trichoderma hfb4 and hfb10 double deletion mutants (N=2 for each genotype) generated in this study. Trichoderma strains grown on PDA at 25 °C.

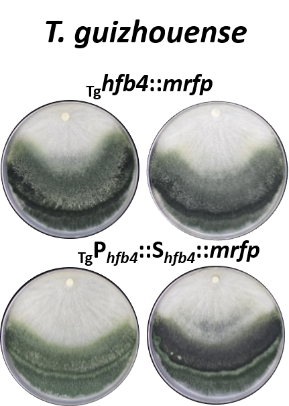

**Figure S5.5** Representative morphology of T. guizhouense mutants expressing mrfp-labelled hfb4 and expressing mrfp under the control of the hfb4 promoter with the hfb4 signal peptide (N=2 for each genotype). Strains grown on PDA at 25 °C for 7 d.

### Supplementary material 6 Intracellular accumulation of mRFP-labelled HFB4 in conidiophores

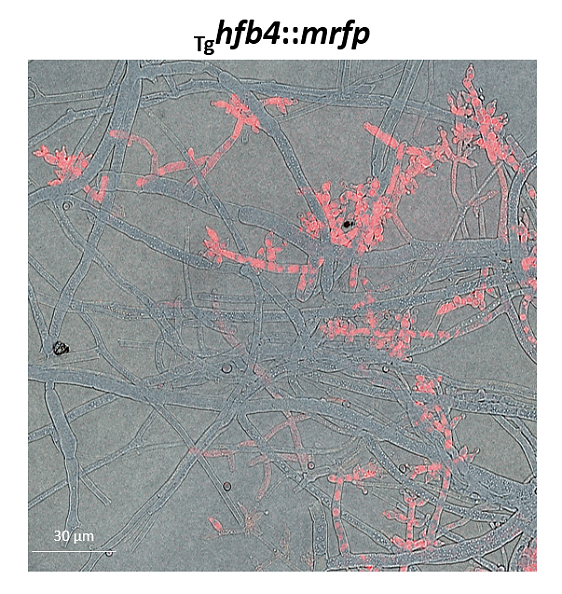

**Figure S6.1** Intracellular accumulation of mRFP-labelled HFB4 in conidiophores of _Tg_*hfb4*::*mrfp* mutant cultivated on cellophane-covered PDA at 25 °C in darkness. The cellophane covered by hyphae was observed in epifluorescent microscope (Leica DMi8 microscope, Germany) without water added.

### Supplementary material 7 Secretion of mRFP expressed using the signal peptide under the control of the promoter of _Tg_*hfb4*

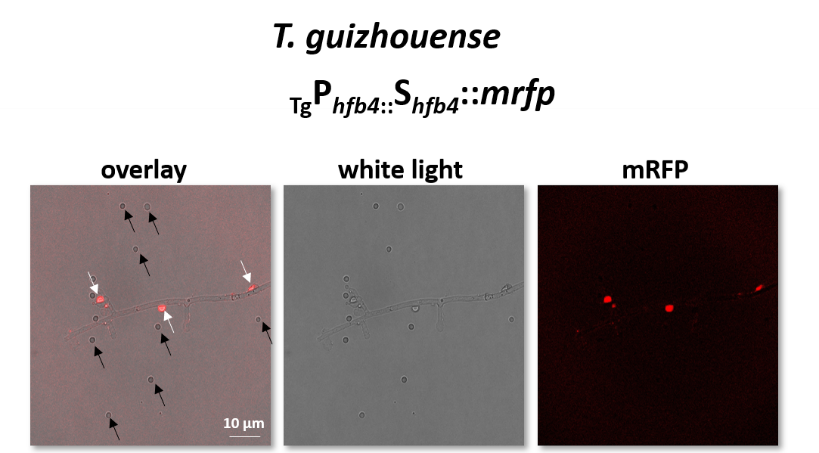

**Figure S7.1** Secretion of mRFP expressed using the signal peptide and the promoter of hfb4 gene from T. guizhouense NJAU 4742. Microscopic analysis of the 48h-old culture grown on PDA at 25 °C in darkness; White arrows point to water/air interface (on air bubbles); black arrows point to spores.

### Supplementary material 8 Representative morphology of *Trichoderma* mutants lacking *rab7*

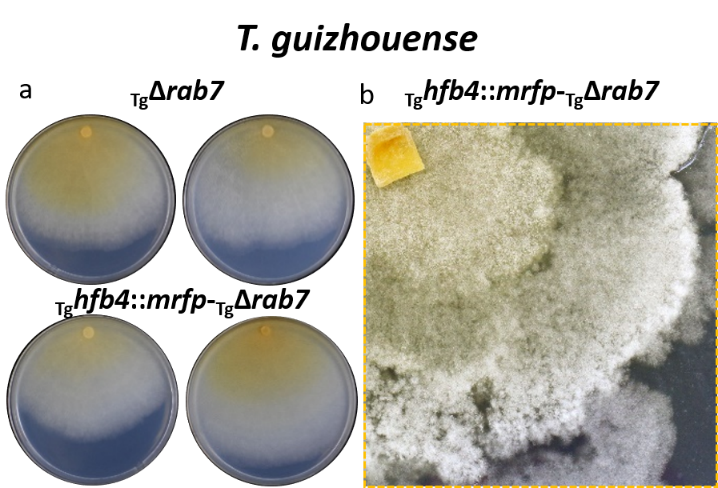

**Figure S8.1** Representative morphology of Trichoderma Δrab7 mutants (N=2 for each genotype) generated in this study (14 d). **a**, Trichoderma strains grown on PDA at 25 °C in darkness; **b**, a close-up image of the colony cultivated at the same conditions for 14 d.

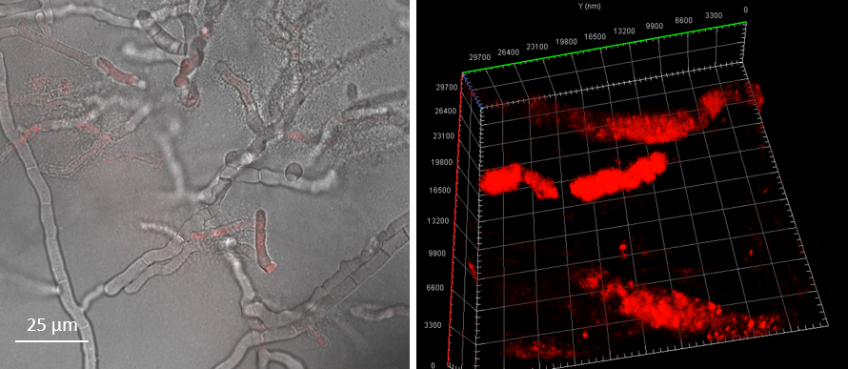

**Figure S8.2** Representative morphology of _Tg_hfb4::mrfp-Δrab7 mutants (N=2 for each genotype observed in epifluorescent microscope when an agar plug was placed of the cover glass. A fluorescence overlay image is sown on the left and a 3D reconstruction of the super-resolution CLSM image is shown on the right.

### Supplementary material 9 Morphology of _Tg_OE*hfb2*::*mrfp* and immunochemical characterization of HFB2 secreted by aerial hyphae

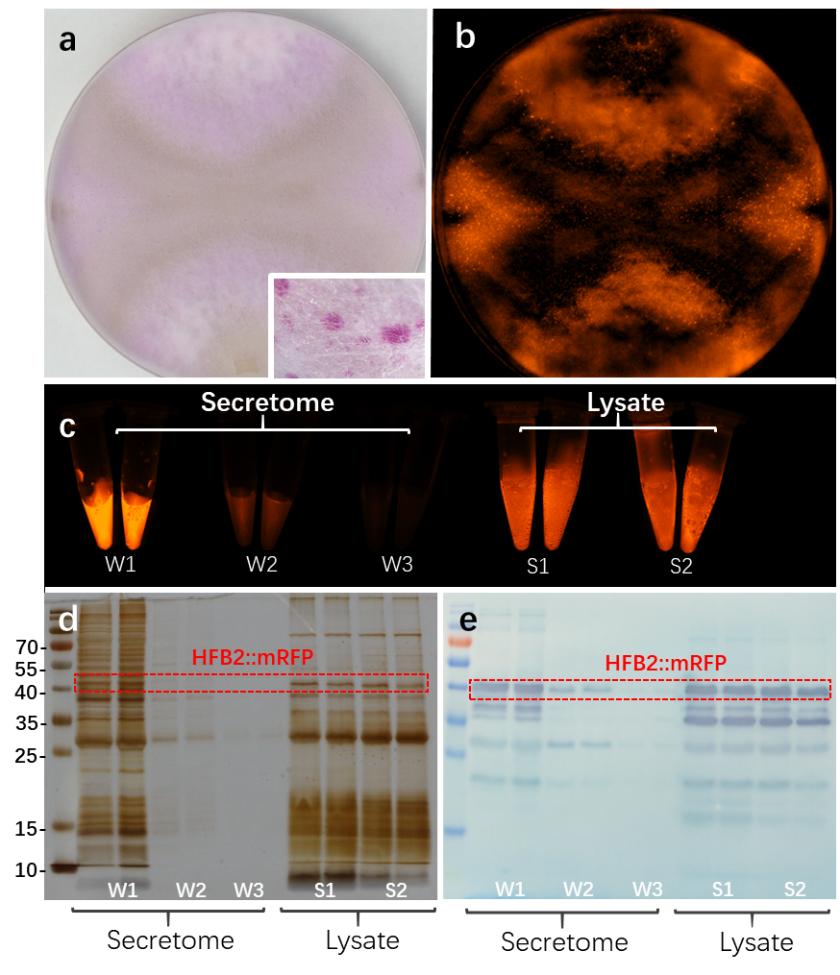

**Figure S9.1** Immunochemical characterization of HFB2 secreted by aerial hyphae of the “pink strain” _Tg_OE*hfb2*::*mrfp*. **a**, morphology of the strain after three weeks of incubation at 25 °C in darkness. The insert in **a** shows a close up view of the aerial hyphae. Note: the aerial mycelium of the wild type *T. guizhouense* strain lives up to 14 -17 d and gets autolyzed. **b**, the same plate imaged by a Bio-Rad ChemiDoc MP (Bio-Rad, USA) equipped with multiplex fluorescent channels. Bright dots correspond to guttation droplets filled with HFB fusion proteins seen in the insert in **a**. **c**, qualitative detection of fluorescence (of HFB2::mRFP) in samples collected from the secretome and the lysate of culture that shown in **a**. The secretome samples (W1-W3) were collected by washing the 21-d-old PDA culture three times with HPLC water. And the lysate samples (S1-S2) were prepared by lysing the water-washed aerial hyphae two times by 1% SDS (dissolved in 50 mM Tris-HCl, pH8.1). **d** and **e**, SDS-PAGE and immune blotting (WB) confirmation of the protein samples shown in **c**.

### Supplementary material 10 Intracellular localization of HFB2::mRFP in two-day-old aerial hyphae of the _Tg_OE*hfb2*::*mrfp* strain

(shown in a separate video file)

### Supplementary material 11 HFB intracellular accumulation in *P. pastoris* strains

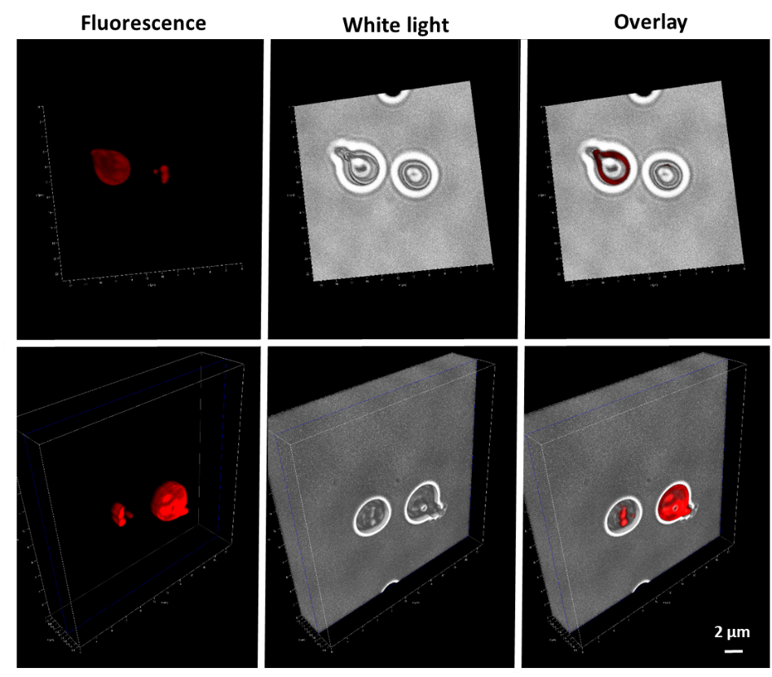

**Figure S11.1** 3D visualization of HFB intracellular accumulation in *P. pastoris* strain P_aox1_S_α_::_Tg_*hfb2*::*mrfp* overexpressing _Tg_*hfb2* was imaged using the Leica DMi8 microscope (Leica, Germany). The strain was cultivated in BMM medium for HFB induction by 0.5 % methanol (see the detailed procedure in Material & Method). An mRFP was fused to the C-terminus of _Tg_HFB2 for protein localization.

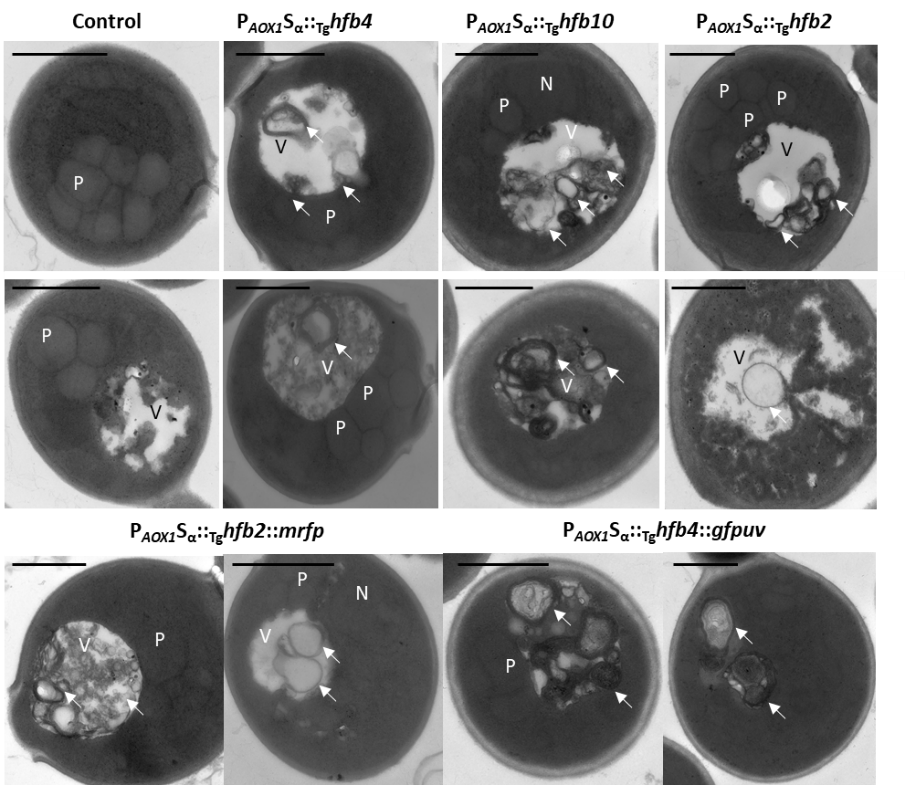

**Figure S11.2** TEM micrographs of the *P. pastoris* mutants overexpressing different HFBs. The control corresponds to the strain transformed with the HFB-free vector (pPICZαA); see the Materials and Methods for the details. P – peroxisome, V – vacuole, N – nucleus. Arrows point to putative HFB vesicles in VMSs that correspond to HFB-enriched organelles. Representative images were selected from total 139 images. Samples for TEM were prepared with at least two mutants and 15 images studied per each haplotype.

### Supplementary material 12 Quantification of autophagy-related genes in *Trichoderma* spp. and *P. pastoris* expressing *hfb*s

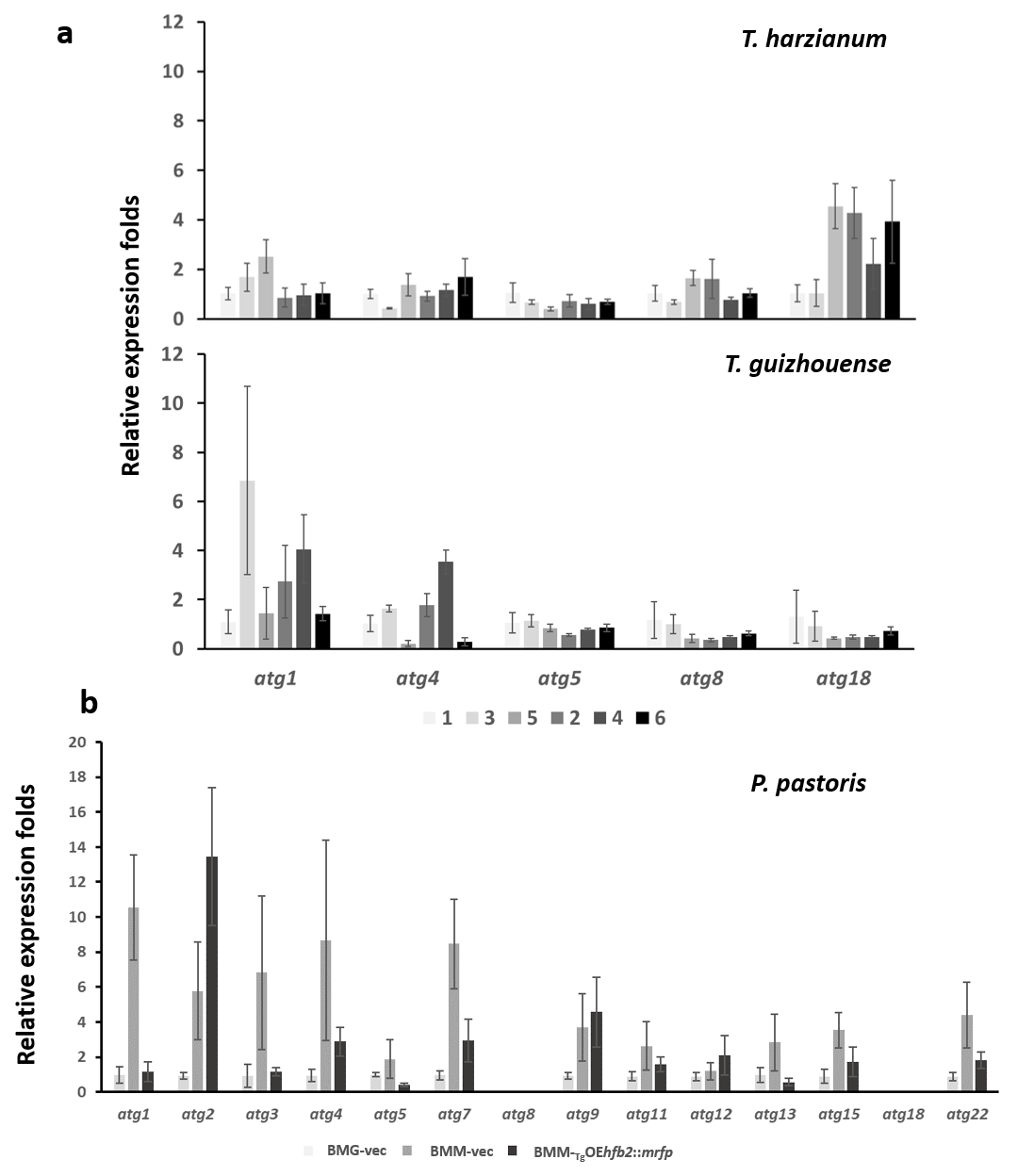

**Figure S12.1** Expression pattern of autophagy-related genes (atg) in Trichoderma spp. during formation and maturation of conidiogenic rings, and in Pichia pastoris strains producing/non-producing HFBs from Trichoderma. The expression ratio of each gene is the mean fold change relative to its expression at the position 1 (see Figure 5) and calibrated by the internal control gene tef1 for Trichoderma spp., and the expression ratio of each gene was compared to its expression in a non-inducible sample (BMG-vec) and calibrated by the internal control gene act1 for P. pastoris, using the ^2-ΔΔ^Ct method. Error bars represent the standard deviation calculated from four biological replicates. RNAs of Trichoderma spp. were extracted from the fungal biomass grown on the desired six positions on the PDA cultures (shown in Figure 5). Position 3 and 4 were the conidiating ring areas. RNAs of P. pastoris were extracted from the yeast cells grown in inducible (BMM)/non-inducible (BMG) cultures. The tested genes were selected based on their reported functions involved in cell autophagy^7, 8, 9^.

### Supplementary material 13 Cell wall ultrastructure of HFB mutants of *Trichoderma*

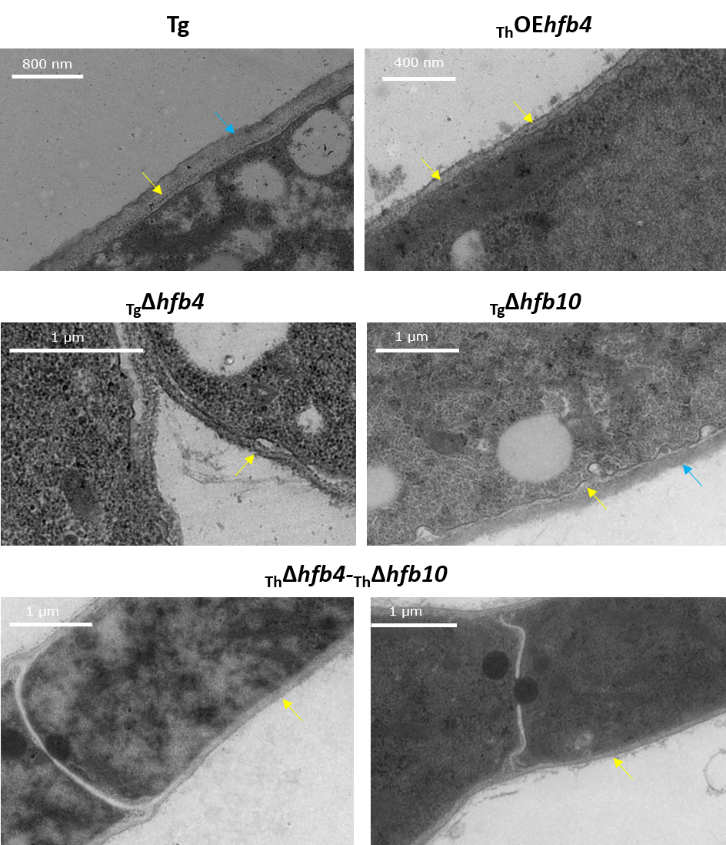

**Figure S13.1** Cell wall ultrastructure of HFB mutants of Trichoderma. Strains cultivated on PDA plates at 25 °C in darkness for 48 h. Yellow arrows point to cell wall, blue arrows indicate the extracellular matrix.

### Supplementary material 14 Colony architecture and dynamic release of HFBs during conidiogenesis of *Trichoderma*

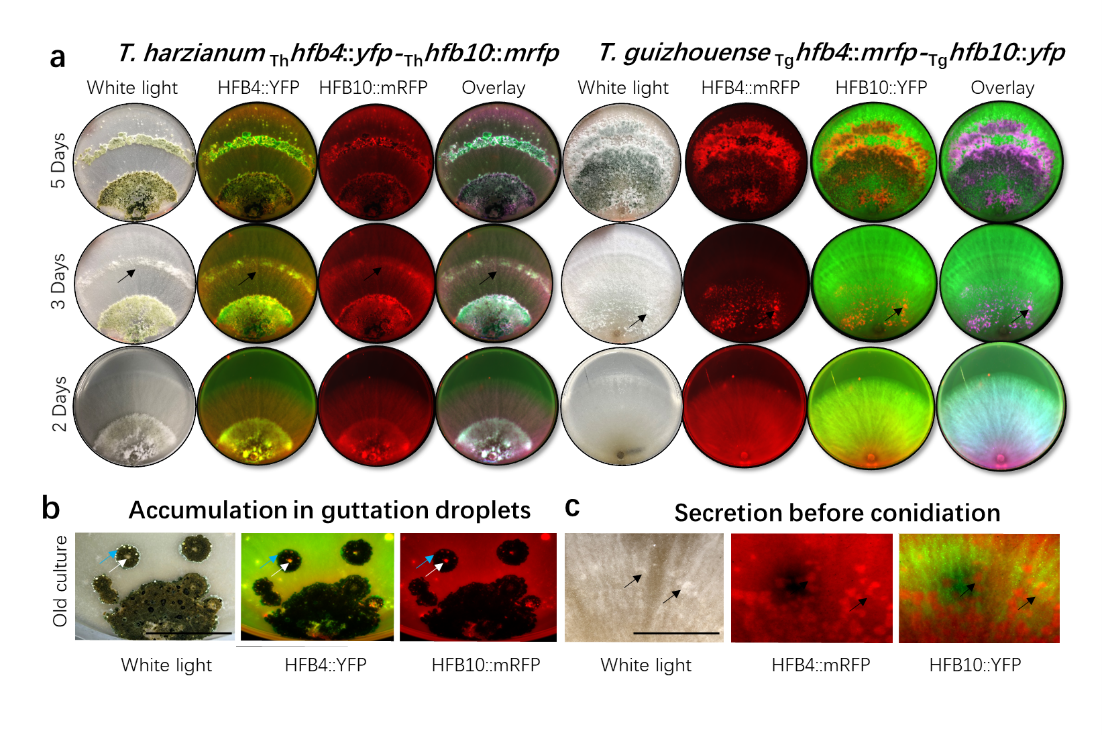

**Figure S14.1** Colony architecture and dynamic release of HFBs by aerial hyphae associated with conidiogenesis. a, Time course-dependent production of HFB4 and HFB10 during the development of *Trichoderma* colonies imaged by a ChemiDoc MP system without magnification. Black arrows point to primordial pustules with premature conidiophores. Note: *T. guizhouense* is autofluorescent in the green spectrum. b, Close-up images obtained from old cultures showing accumulation of HFBs in guttation droplets above conidia (white arrows) but not in drops above young hyphae (blue arrows). c, Production of HFB4 and HFB10 at the spots of subsequent formation of conidiophores and spores. Primordial pustules are shown by black arrows.

### Supplementary material 15 Animated 3D reconstructions of extracellular HFB-enriched matrices coating sporulating *Trichoderma* colonies

(shown in a separate video file)

### Supplementary material 16 Video showing putative HFB-enriched vesicles secreted by _Tg_OE*hfb2*::*mrfp* strain

(shown in a separate video file)

### Supplementary material 17 Supplementary results and detailed methods

#### S17.1 Gene deletion

HFB-encoding genes in genomes of *T. harzianum* CBS 226.95 (GenBank: MBGI00000000.1) and *T. guizhouense* NJAU 4742 (GenBank: LVVK00000000.1) were obtained by genome mining as given in our previous work^10^ and gene IDs were listed in **Supplementary material 1**. Vectors for gene deletion was constructed as Cai et al.^10^ described. Briefly, gene of interest was replaced by *hph* cassette or by *neo* cassette by transformation. Positive mutants were purified by the method of single spore isolation and confirmed for the absence of the target gene with the primer pair F3 and R3. All vectors and PCR products were confirmed by sequencing; similarly hereinafter. After purification and verification by PCR (shown in **Figure S17.1** with two randomly selected mutants).

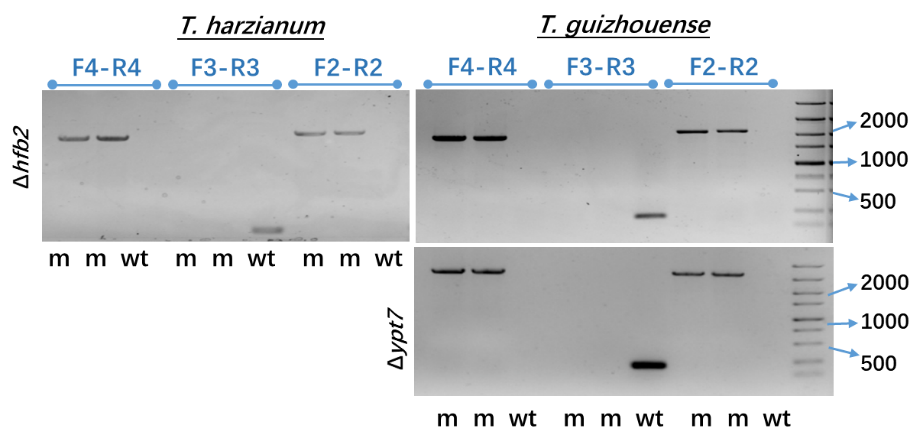

**Figure S1** PCR verification of gene deletion mutants. wt represents the corresponding wild type strains; m represents mutants. DL5000 DNA marker (Vazyme, China) was used in gel electrophoresis

#### S17.2 *in situ* fluorescence labelling

Schematic diagram for vector construction of fluorescent labelling were shown in **Figure S17.2**. Fragments were respectively amplified by PCR and fused into PUC19 together with the selected fluorescent protein gene and the selection marker cassette. A flexible linker sequence (GGGGS×3)^11^ was designed to connect the HFB protein and the fluorescent protein. And a 6×His tag was fused to the C-terminus of the fluorescent protein, which allows an immunological detection for the fused protein. Specifically, HFB4 in *T. guizhouense* NJAU 4742 strain was labelled by mRFP using *hph* as the marker and then HFB10 was labelled by YFP using *neo* as the marker. In *T. harzianum* CBS 226.95, HFB4 was labelled by YFP with *hph* and HFB10 was with mRFP and *neo*, in order to rule out the position effect of the labelling sequence on protein localization. The hph cassette, neo cassette and mrfp gene were cloned from plasmids of pPcdna1-hph, pKi-Gen and pPICZα-mRFP that were maintained in Microbiology and Comparative Genomics group of TU Wien (Austria). The *yfp* gene was cloned from the plasmid of pDS22 that was kindly provided by Dr. Norio Takeshita and Dr. Reinhard Fischer (Institute for Applied Biosciences, Karlsruhe Institute of Technology, Germany). After purification, one double-labelled mutant was generated for each species. And the results of *hfb* labelling were shown in **Figure S17.3**.

Additionally, a mutant expressing mRFP after the signal peptide of *hfb4* under the control of the native promoter of *hfb4* (P*_hfb4_*) was generated (**Figure S17.4**). A 1.2 kb fragment containing the *hfb4* promoter in frame with the HFB4 native signal peptide sequence (i.e., ATGAAGTTCTCTGCCATCGCTCTCTTCGCCTCGCTGGCCATTGCCGCGCCCGCCACGGAGGCC) and a 1.2 kb fragment containing the native transcription termination of *hfb4* were cloned from *T. guizhouense* NJAU 4742 and fused into PUC19 plasmid together with mRFP gene and the selection marker cassette. The vector was constructed to allow the expression of mRFP under the control of P*_hfb4_* and its secretion led by the secretion signal of HFB4. One transformant was screened out harboring the correct construct (shown in **Figure S17.5**).

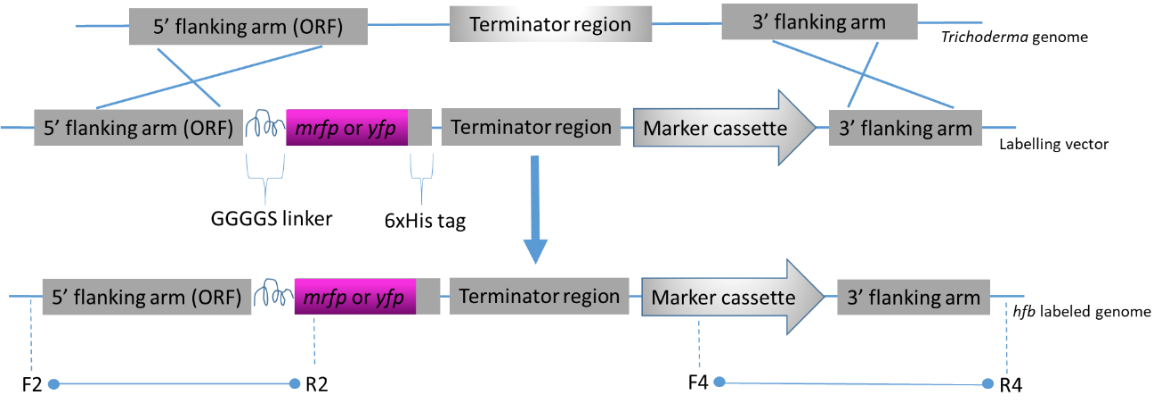

**Figure S17.2** Schematic diagram of hfb labelling via homologous recombination and mutant screening

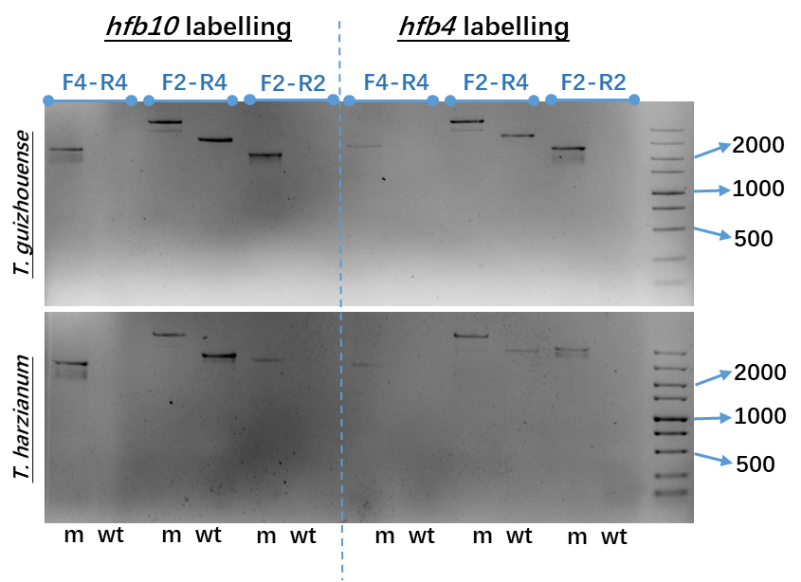

**Figure S17.3** PCR verification of the double-labelled mutants. wt represents the corresponding wild type strains; m represents mutants. DL5000 DNA marker (Vazyme, China) was used in gel electrophoresis.

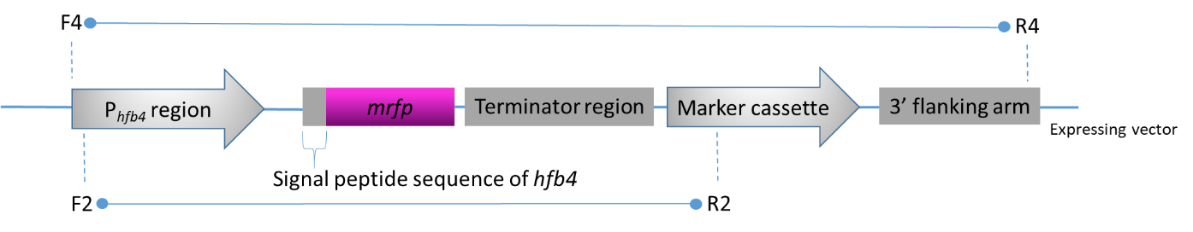

**Figure S17.4** Schematic diagram of expressing mRFP under the native promoter of hfb4 (P_hfb4_) from T. guizhouense NJAU 4742 and mutant screening.

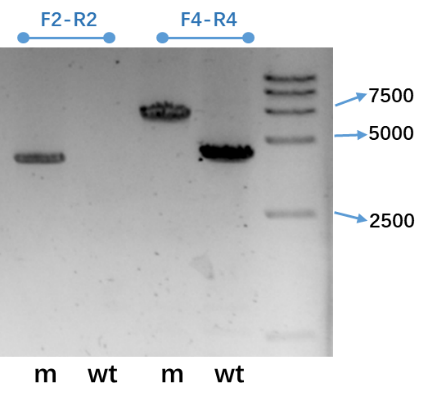

**Figure S17.5** PCR verification of mutant expressing mRFP. wt represents the wild type strain of T. guizhouense NJAU 4742; m represents mutants. DL15000 DNA marker (Vazyme, China) was used in gel electrophoresis.

#### S17.3 Overexpression of *hfb* encoding genes in *Trichoderma*

Overexpressing vectors were constructed as shown in **Figure S17.6**. The open reading frame (ORF) of *hfb* gene and its native terminator region was cloned from *Trichoderma* genomic DNA and inserted into the pUCPcdna1-hph plasmid (Cla I pre-digested) after a constitutive promoter P*_cdna1_* from *T. reesei* QM6a^12^. After purification and verification by PCR (shown in **Figure S17.7** with two randomly selected mutants), five _Th_OE*hfb4* and two _Th_OE*hfb10* were obtained for *T. harzianum* CBS 226.95 and three _Tg_OE*hfb4* and two _Tg_OE*hfb10* mutants were obtained for *T. guizhouense* NJAU 4742.

For overexpressing *hfb2* with fluorescent tag (*mrfp*) in *T. guizhouense* NJAU 4742 under the constitutive promoter P*_cdna1_*, a 0.5 kb fragment of *hfb2* containing the ORF and a 1.3 kb terminator region from the genomic DNA of *T. guizhouense* NJAU 4742 were obtained by PCR. The PCR products were purified and fused together with *mrfp* gene into ClaI-digested pPcdna1-hph plasmid as the order shown in **Figure S17.8**. A GGGGS ×3 linker and a His ×6 tag were introduced into the construction during the primer synthesis as mentioned above. The transformation resulted into two positive mutants confirmed by PCR shown in **Figure S17.9**.

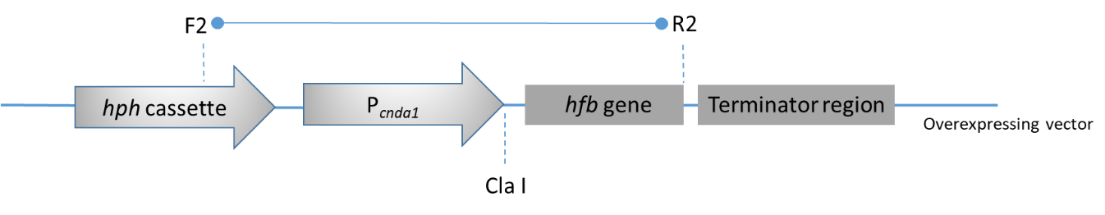

**Figure S17.6** Schematic diagram of overexpressing hfbs under a constitutive promoter P_cnda1_ and mutant screening.

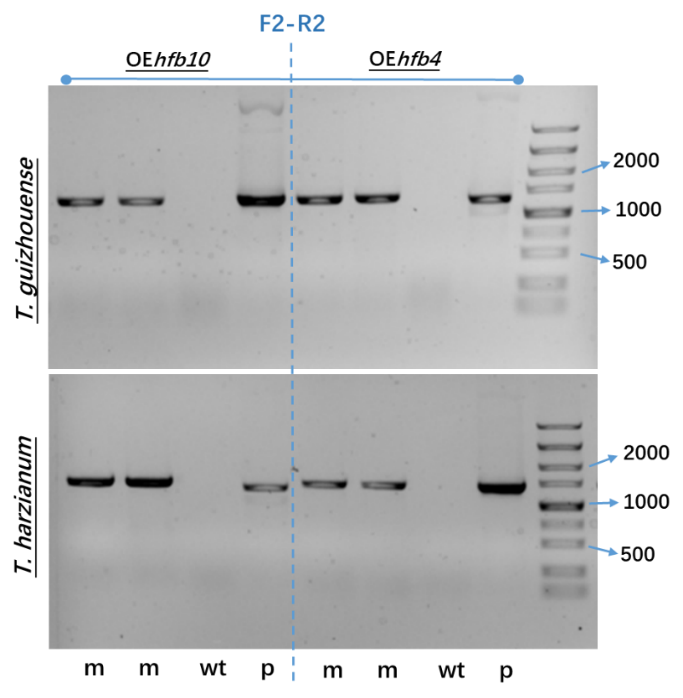

**Figure S17.7** PCR verification of mutant overexpressing hfb4 or hfb10. wt represents the corresponding wild type strains; m represents mutants; p, positive control cloned from the corresponding plasmid. DL5000 DNA marker (Vazyme, China) was used in gel electrophoresis.

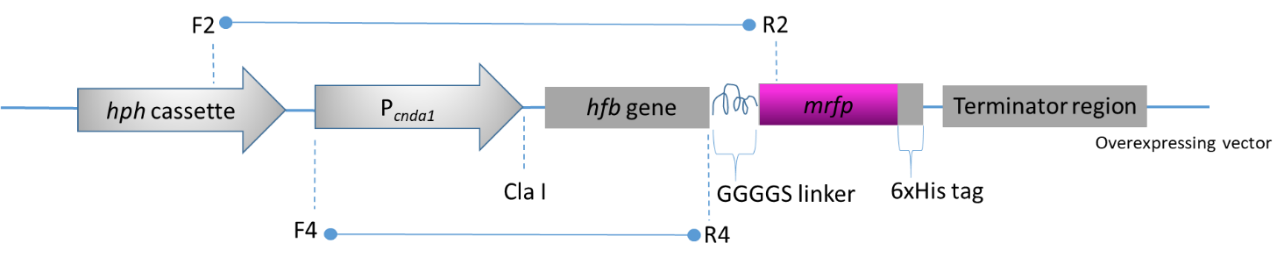

**Figure S17.8** Schematic diagram of overexpressing mrfp-fused hfbs under a constitutive promoter P_cnda1_ and mutant screening.

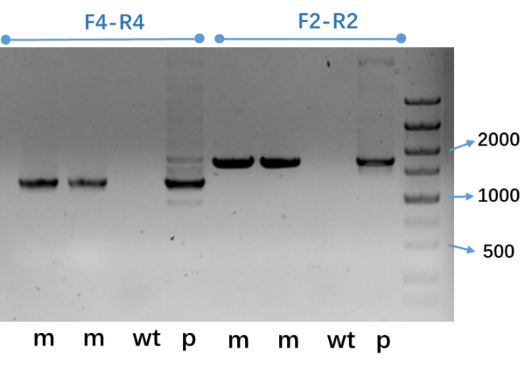

**Figure S17.9** PCR verification of mutant overexpressing mrfp-fused hfb2 in T. guizhouense NJAU 4742. wt represents the corresponding wild type strains; m represents mutants; p, positive control cloned from the corresponding plasmid. DL5000 DNA marker (Vazyme, China) was used in gel electrophoresis.

#### S17.4 Reverse complement of *hfb* encoding genes in *hfb*-deletion mutants

Reverse complement vectors were constructed as shown in **Figure S17.10** with two strategies, namely with or without the fluorescent tag. The open reading frame (ORF) of *hfb* gene and its native promoter and terminator region was amplified from *Trichoderma* genomic DNA and fused with the *neo* cassette. After purification and verification, PCR confirmation of each genotype is shown in **Figure S17.11** with two randomly selected mutants.

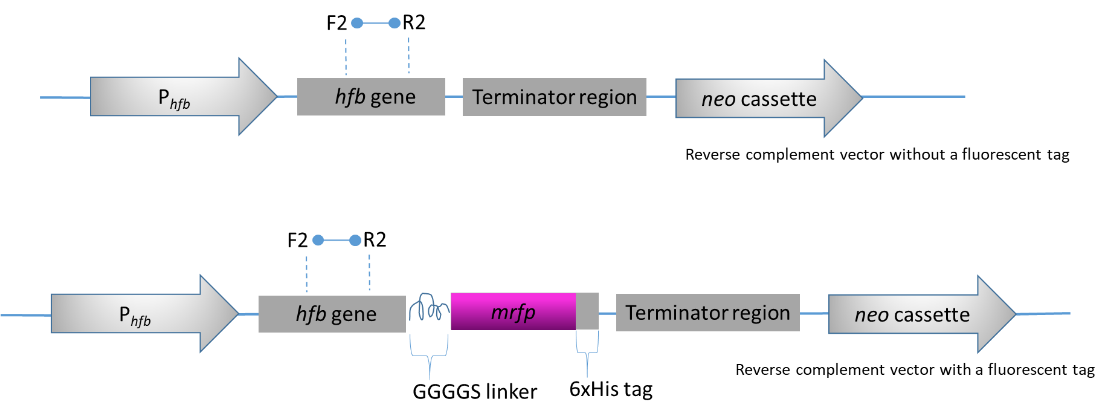

**Figure S17.10** Schematic diagram of reverse complement of hfbs to the respective hfb-deletion mutant.

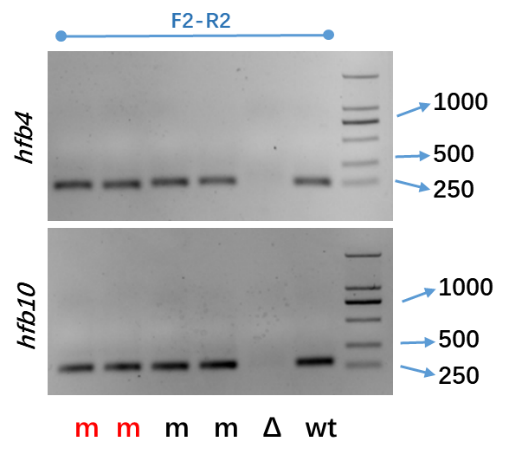

**Figure S17.11** Transcriptional verification (RT-PCR) of mutant reverse complemented with hfb4 or hfb10 with or without a fluorescent tag. wt represents the corresponding wild type strains; m represents mutants (red font highlights the fluorescently labelled ones); Δ, represents the hfb-deletion mutant. DL5000 DNA marker (Vazyme, China) was used in gel electrophoresis. RNA extraction was carried out for the 48-h-old PDA cultures of each genotype, and cDNA synthesis was performed as described in the Materials and Methods.

#### S17.5 Heterologous expression of *hfb* encoding genes from *Trichoderma* in *P. pastoris*

The EasySelect^TM^ *Pichia* Expression Kit was used to express genes from *Trichoderma* in *P. pastoris* strain KM71H, according to the manufacturer’s instructions (Invitrogen, USA). The amplified *hfb* gene (without signal peptide or intron sequences) was inserted into the position between the restriction site of EcoR I and Xba I of plasmid pPICZαA (**Figure S17.12**). To express recombinant proteins with a fluorescent tag, the GFPuv or the mRFP was adopted and fused at the C-terminus of the *hfb* (**Figure S17.13**). Besides, the native Saccharomyces cerevisiae α-factor secretion signal was synthesized at the N-terminus of the *hfb* gene and a His ×6 epitope at the C-terminus. Zeocin resistance driven by the *sh ble* cassette was used for selection. The electroporation resulted in five _Tg_OE*hfb4*, four _Tg_OE*hfb10*, ten _Tg_OE*hfb2*, five _Tg_OE*hfb4-gfpuv* and six _Tg_OE*hfb2-mrfp* were obtained (shown in **Figure S17.14** with one randomly selected mutant).

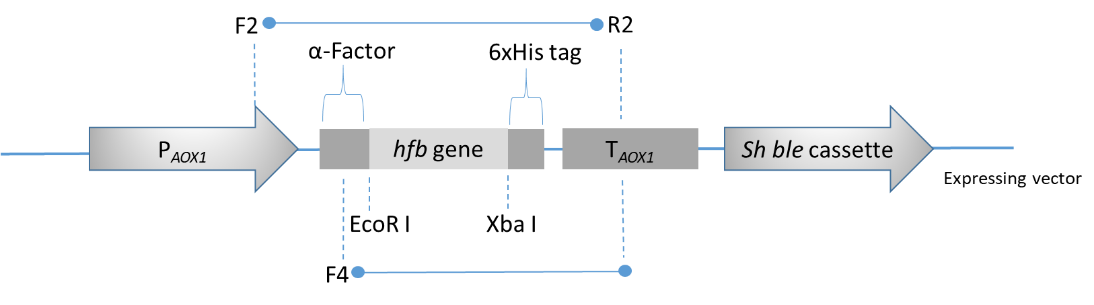

**Figure S17.12** Schematic diagram of expressing hfbs under a methanol-inducible promoter P_AOX1_ in P. pastoris and mutant screening. α-Factor, native S. cerevisiae α-factor secretion signal; T_AOX1_, native transcription termination from AOX1 gene of P. pastoris; Sh ble cassette, from Streptoalloteichus hindustanus ble gene driving resistance to Zeocin.

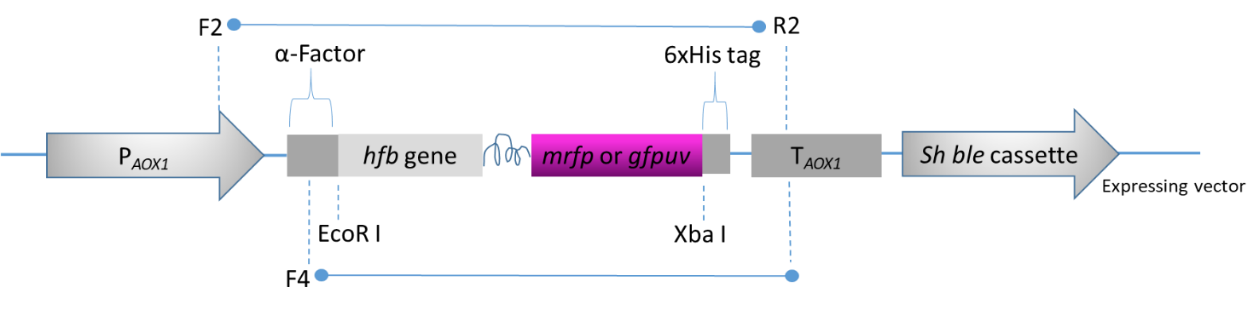

**Figure S17.13** Schematic diagram of expressing fluorescently tagged hfbs under a methanol-inducible promoter P_AOX1_ in P. pastoris and mutant screening. α-Factor, native S. cerevisiae α-factor secretion signal; T_AOX1_, native transcription termination from AOX1 gene of P. pastoris; Sh ble cassette, from S. hindustanus ble gene driving resistance to Zeocin.

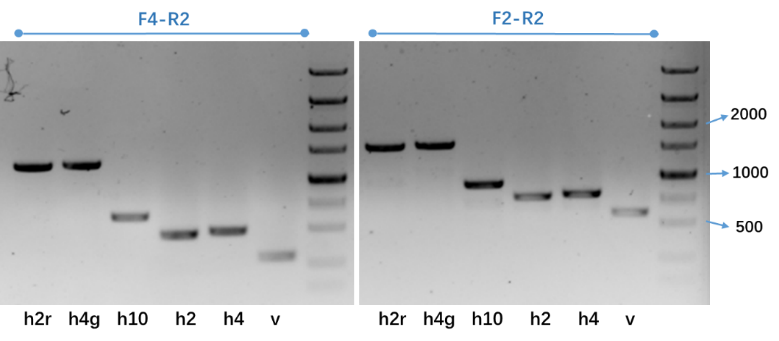

**Figure S17.14** PCR verification of mutant overexpressing hfbs (from T. guizhouense NJAU 4742) or fluorescently tagged hfbs in P. pastoris. h4 represents mutants harboring hfb4; h2 represents mutants harboring hfb2; h10 represents mutants harboring hfb10; h4g represents mutants harboring gfpuv-fused hfb4; h2r represents mutants harboring mrfp-fused hfb2; v represents mutants transformed with the original vector pPICZαA without hfbs. DL5000 DNA marker (Vazyme, China) was used in gel electrophoresis.

#### S17.6 Immunoblotting assays

For Western blotting (WB), the protein samples obtained from the PDA culture of labelled mutants and their corresponding wild type strains were separated by SDS-PAGE and then were transferred to nitrocellulose membrane (GE Healthcare) at 100 V for 1 h using Mini Trans Blot Cell (Bio-Rad, USA) with wet transfer technique. The membrane was treated with Clarity^TM^ Western ECL and imaged by the Bio-Rad ChemiDoc MP system (Bio-Rad, USA) or treated with the ONE-HOUR Western^TM^ Standard Kit (Genscript, China) and imaged by Cannon EOS 70D, according to manufacturer’s instructions. HRP conjugated Anti-His Tag Mouse Monoclonal antibody (Invitrogen, USA) was used at 1:2000 dilution. The WB results shown in **Figure S17.15** gave the corrected size (ca. 40 KDa) of bands of fused HFB::mRFP or HFB::YFP proteins in *Trichoderma*. The SDS-PAGE (silver stained) and WB results shown in **Figure S17.16** demonstrated the expression of HFBs in *P. pastoris*. The extra bands with lower molecular weights may be due to the cleavage of the proteins.

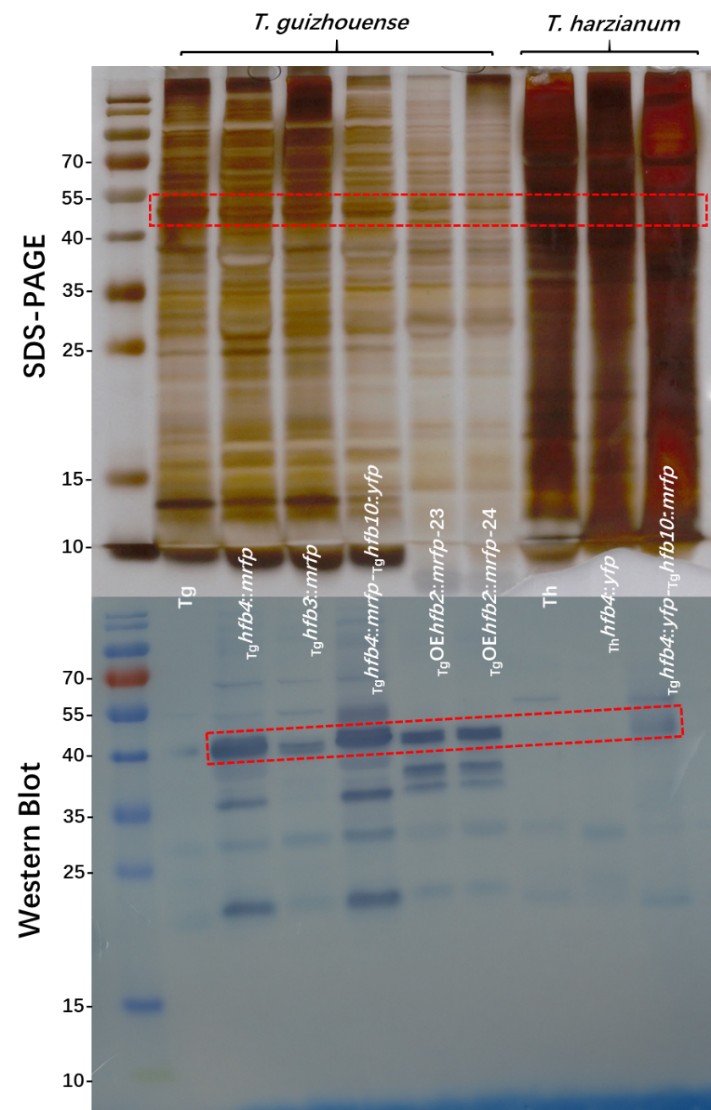

**Figure S17.15** Western Blot confirmation of the labelled strains of Trichoderma (WB visualization was performed by the ONE-HOUR Western^TM^ Standard Kit, Genscript, China). Protein samples were collected by washing the 5-day-old PDA culture with HPLC water. Bands framed represent the target bands ca. 40 KDa. PageRuler™ Prestained Protein Ladder (Fermentas, USA) was used in gel electrophoresis.

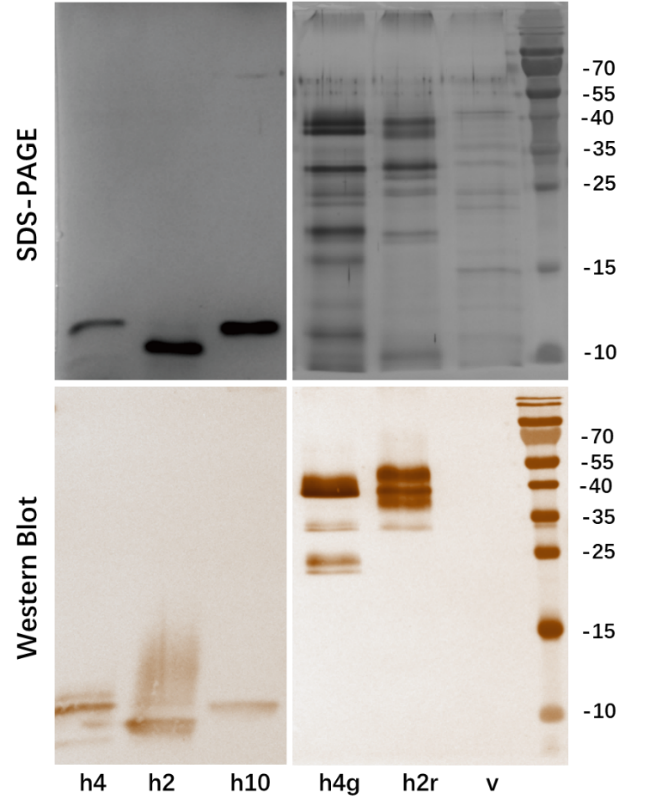

**Figure S17.16** Western Blot confirmation of the labelled strains (performed by Clarity^TM^Western ECL, Bio-Rad, USA). h4 represents mutants overexpressing hfb4; h2 represents mutants overexpressing hfb2; h10 represents mutants overexpressing hfb10; h4g represents mutants overexpressing gfpuv-fused hfb4; h2r represents mutants overexpressing mrfp-fused hfb2; v represents mutants transformed with the original vector pPICZαA without hfbs.

### Supplementary material 18 Primers used in this study

(shown in a separate file)
